# Structural basis for stereospecific inhibition of ASCT2 from rational design

**DOI:** 10.1101/2020.05.29.124305

**Authors:** Rachel-Ann A. Garibsingh, Elias Ndaru, Alisa A. Garaeva, Massimiliano Bonomi, Dirk J. Slotboom, Cristina Paulino, Christof Grewer, Avner Schlessinger

## Abstract

ASCT2 (SLC1A5) is a sodium-dependent neutral amino acid transporter that controls amino acid homeostasis in peripheral tissues. ASCT2 is upregulated in cancer, where it modulates intracellular glutamine levels, fueling cell proliferation. Nutrient deprivation via ASCT2 inhibition provides an emerging strategy for cancer therapy. Here, guided by a homology model of ASCT2 in an outward-facing conformation, we rationally designed novel inhibitors exploiting stereospecific pockets in the substrate binding site. A cryo-EM structure of ASCT2 in complex with inhibitor (*Lc*-BPE) validated our predictions and was subsequently refined based on computational analysis. The final structures, combined with MD simulations, show that the inhibitor samples multiple conformations in the ASCT2 binding site. Our results demonstrate the utility of combining computational modeling and cryo-EM for SLC ligand discovery, and a viable strategy for structure determination of druggable conformational states for challenging membrane protein targets.

---

The <u>A</u>lanine-<u>S</u>erine-<u>C</u>ysteine <u>T</u>ransporter 2 (SLC1A5, ASCT2) is a sodium-dependent transporter of neutral amino acids that is expressed at low levels in various tissues including the intestine, kidney, liver, heart, placenta, and brain^1,2^. ASCT2 belongs to SLC1 family, which in humans includes glutamate transporters EAAT1-5 and neutral amino acid transporters ASCT1-2. ASCT2 is highly upregulated in several tumor types, such as triple negative breast cancer (TNBC), prostate cancer, and melanoma (reviewed in reference 3), where it imports glutamine into cells that is utilized to build biomass and enhance proliferation *via* mTORC1^4^, a process driven by the transcription factor c-MYC^5^. Recently, similar processes have been identified in the differentiation and activation of T-cells, as well as in programmed cell^6,7^, expanding the relevance of ASCT2 to central nervous system (CNS) disorders, heart disease, and other conditions. Notably, inhibition of ASCT2 has been shown to reduce intracellular glutamine levels and subsequently tumor size *in vivo*^*8*^. Taken together, ASCT2 is emerging as an important therapeutic target for a multitude of diseases and disorders.

A prerequisite for developing clinically-relevant inhibitors for ASCT2, is the structural characterization of ASCT2’s distinct conformational states, as well as its mode of interaction with inhibitors. Multiple structures of human SLC1 members and their prokaryotic homologues, combined with biophysical data, have demonstrated that these transporters operate using an elevator transport mechanism^9-14^. In this mechanism, the transport domain with bound substrate moves perpendicularly to the membrane, while the static scaffold domain remains in place and constitutes the trimeric subunit interface. The conservation of the structure and mechanism across the SLC1 family has enabled the generation of high-quality models for human SLC1 members, capturing different snapshots of the transport cycle and aided the development of novel inhibitors^15-18^.

Recently determined cryo-EM structures of human ASCT2 in various conformations have visualized distinct steps of the ASCT2 transport cycle for the first time^19,20^. Specifically, the inward-occluded structure of ASCT2 with bound glutamine was solved first^20^, confirming the hypotheses proposed based on homolog structures, particularly in terms of fold and transport mechanism. Subsequently, the inward-open structure of ASCT2 suggested that the transporter works using a single gate mechanism, where hairpin 2 (HP2) controls access to the substrate binding site on both the intracellular and extracellular sides of the membrane. The structure also demonstrated that compounds can bind in the previously predicted sub-pocket region in the binding site, called pocket A (Extended Data Fig. 1)^19^. Both the inward-open and outward-occluded structures of ASCT2 showed a putative cholesterol binding site at the interface between mobile and scaffold domains^19,21^. Importantly, the ASCT2 structures are consistent with previous functional observations based on homology models, increasing the confidence in the computational methods used to study this family of proteins.

Because ASCT2 is a pharmacologically important target, multiple small molecule competitive inhibitors have been developed for this protein, using both structure-based rational design and ligand-based approaches (reviewed in reference 22). For example, guided by homology modeling, we discovered and optimized a series of proline-based derivatives as well as other chemical scaffolds that inhibit ASCT2 in low µM potencies^16,23,24^. Notably, these amino acid analog inhibitors, which have large hydrophobic groups in the side chain provided evidence that non-polar interactions within the substrate binding site may be exploited to further increase potency. However, the low potency and selectivity of these inhibitors, combined with the lack of high resolution confirmation of their binding mode *via* experimentally solved structures, have hindered optimization efforts to develop clinically relevant compounds. Additionally, other human SLC1 members, EAAT1 (SLC1A3)^25^ and EAAT2 (SLC1A2)^26^ have been targeted by non-competitive allosteric modulators, suggesting that similar strategies could be applied to ASCT2. A potential limitation of such compounds is their increased lipophilicity and low solubility, hindering their utility in potential clinical applications.

Here, we describe an integrated approach using computer-aided compound design, functional testing, and structure determination with cryo-EM, to rationally design novel inhibitors for ASCT2. We first refined an outward-facing homology model of ASCT2^27^, and used molecular docking, electrophysiology, and transport assays in proteoliposomes to develop and characterize novel and potent stereoselective ASCT2 inhibitors targeting a druggable sub-pocket in the substrate binding site. Next, we determined the cryo-EM structure of ASCT2 in complex with the most potent inhibitor: L-*cis* hydroxyproline biphenyl ester (*Lc*-BPE). Subsequently, using iterative homology model refinement in combination with molecular dynamics (MD) simulations with the cryo-EM density map as a restraint, we characterized pharmacologically relevant conformations of the inhibitor in the structure. Finally, we discuss the relevance of our approach to describe the structural basis of ASCT2 inhibition, as well as its utility in guiding structure determination of challenging SLC targets in pharmacologically relevant conformations.

## RESULTS

### Overview of the approach

We began by iteratively refining the binding site of a homology model of ASCT2, based on an outward-open crystal structure of human EAAT1 (EAAT1; 46% sequence identity) (Fig. 1). Each model was evaluated for its ability to distinguish known ligands from decoys or likely non-binders, with ligand docking (i.e., ligand enrichment calculations). Virtual compounds were then docked against the model and top predictions were synthesized and tested using electrophysiology. This approach enabled us to progressively optimize the binding site for protein-ligand complementarity for the rational design of four new ASCT2 inhibitors. Next, we characterized the mechanism of inhibition of the most potent inhibitor (*L*c-BPE) using biochemical and biophysical approaches. A cryo-EM structure of ASCT2 was subsequently determined with bound *L*c-BPE, revealing subtle differences in the ligand binding pose and sidechain placement of binding site residues as compared to the models. Further analysis of the cryo-EM density map and homology model, followed by MD simulations, facilitated the identification of distinct ASCT2 and inhibitor conformations, and demonstrated the flexibility of the HP2 loop.

**Figure 1.**
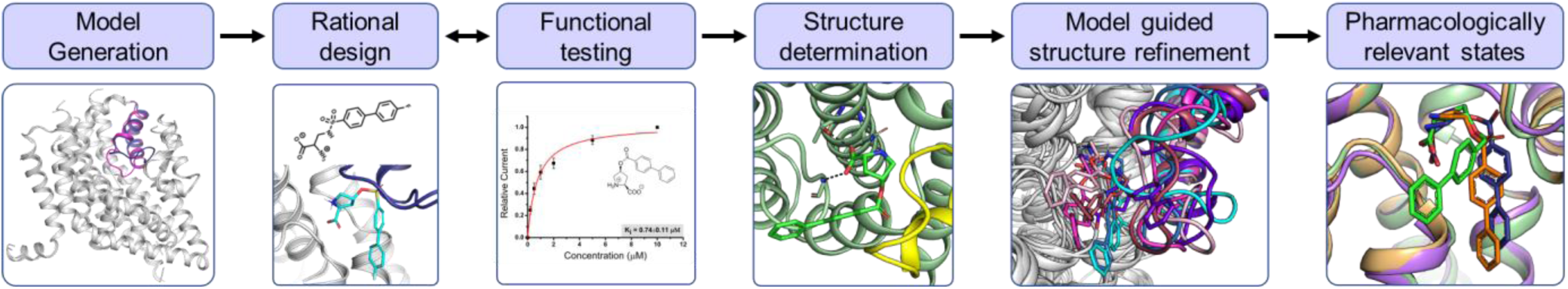
Overview of modeling-guided structural characterization of ASCT2-inhibitor binding. Initial homology model was constructed and refined based on homolog structures. The model is then subjected to rational compound design by iteratively performing molecular docking, chemical synthesis, and functional testing, as well as ligand enrichment calculations. Cryo-EM is then used to determine and confirm the complex structure and is subsequently refined using the homology model and MD simulations, to identify distinct pharmacologically relevant conformations of ASCT2.

### ASCT2 modeling and inhibitor design

We have previously developed a series of ASCT2 inhibitors based on proline sulfonamide and sulfonic acid esters predicted to target a subpocket of the ASCT2 substrate binding site (pocket A), guided by homology models in the outward-open conformation^23^. We found that molecules with a linker allowed compounds to dock into pocket A, and the inhibitor’s sidechain lipophilicity correlated with experimental K_i_ values. Further, the models correctly predicted the importance of a hydrophobic biphenyl group in the sidechain for efficient inhibitor binding^23^. Based on analysis of recent ASCT2 structures in the inward-open and the outward-occluded conformations^15,16,23^, combined with docking of known ligands, we remodeled the sidechains of S354, D464, and C467 (Materials and Methods). The resultant model obtained ligand enrichment score higher than those enrichments of any previously described cryo-EM structures or models of ASCT2 (e.g., AUC of 94.3 as compared to AUC of 75 for the inward open structure), suggesting the model represents an alternative pharmacologically relevant conformation.

Additionally, docking of proline sulfonamide and sulfonic acid esters revealed that *trans* isomers with substitutions to the 4-position of the proline-like scaffold: (i) pointed on opposite sides of the proline ring plane as expected from their *trans* configuration; and (ii) docked less favorably compared to *cis* isomers^23^. We hypothesized that *cis* isomers would force the linker moiety of the proline-like scaffold to point towards the base of pocket A, while maintaining critical polar interactions with hairpin 1 (HP1) and TM8 (e.g. with S353 and N471), typical of ASCT2 substrates and inhibitors^20,21^. Further, to improve the solubility of the inhibitors while maintaining activity on ASCT2, we substituted the sulfonamide and sulfonic acid ester linkages with an ester linker and therefore synthesized four novel ester derivatives based on the 4-hydroxyproline amino acid scaffold. These four proline biphenyl esters (BPEs) are diastereomers with the L- (*L*c-BPE and *Lt*-BPE) and D-configuration (*Dc-*BPE and *Dt*-BPE) at the α-amino acid carbon. We then used molecular docking and estimated the free energy of binding using MM-GBSA, to assess binding of the new ligand series (Material and Methods; Table 1). Interestingly, both docking scores and MM-GBSA calculations of *cis* and *trans* isomers of the L and D compounds (Extended Data Fig. 1, Table 1) suggest that *cis* compounds are more potent than their *trans* counterparts. We also observe that the L-amino acid scaffold is preferred to the D-amino acid, in agreement with previous reports^28^.

**Table 1:**
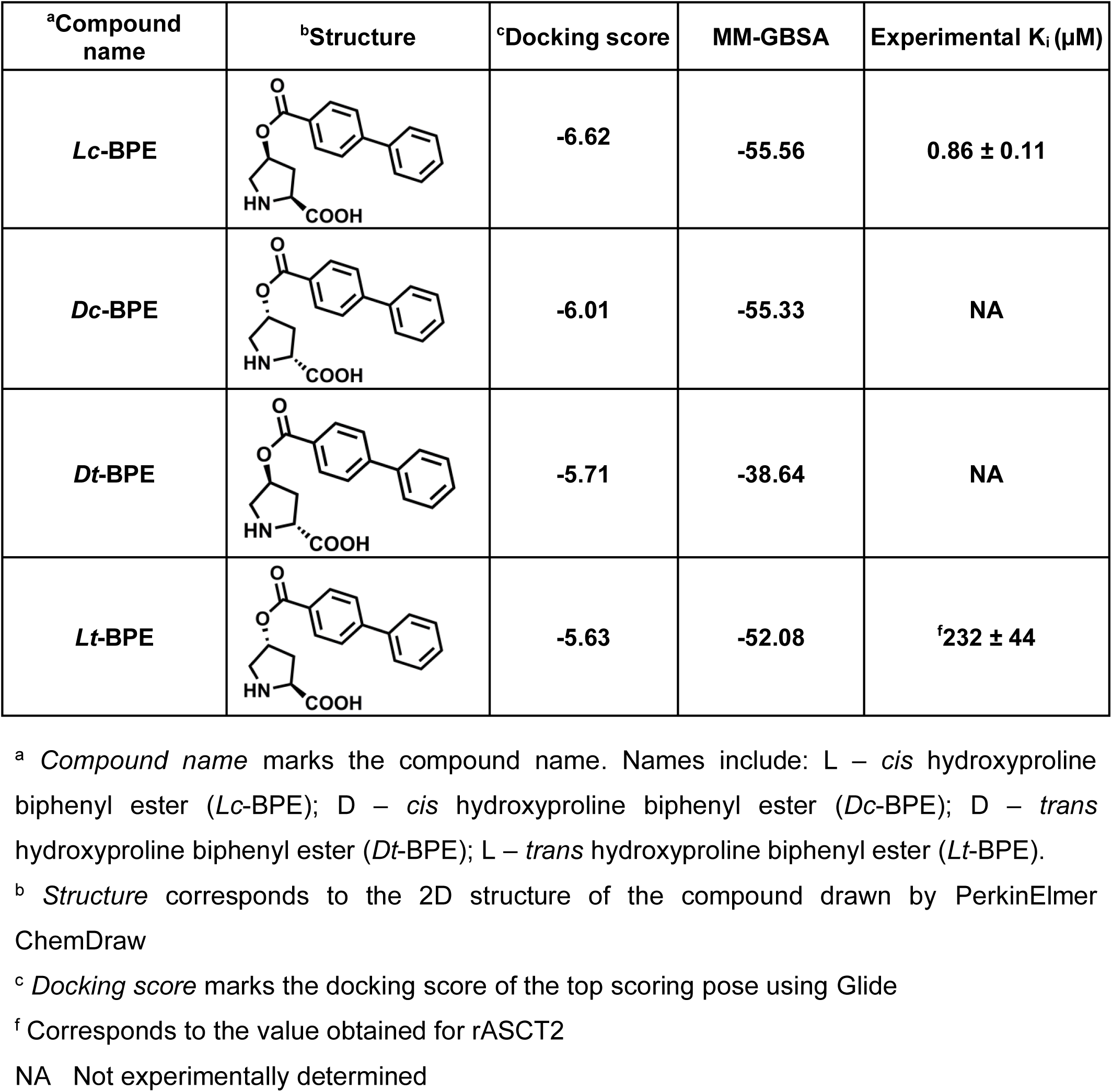
Tool compounds developed in this study.

| <sup>a</sup>Compound name | <sup>b</sup>Structure | <sup>c</sup>Docking score | MM-GBSA | Experimental K<sub>i</sub> (μM) |
| --- | --- | --- | --- | --- |
| *Lc*-BPE |  | -6.62 | -55.56 | 0.86 ± 0.11 |
| *Dc*-BPE |  | -6.01 | -55.33 | NA |
| *Dt*-BPE |  | -5.71 | -38.64 | NA |
| *Lt*-BPE |  | -5.63 | -52.08 | <sup>f</sup>232 ± 44 |

<sup>a</sup> *Compound name* marks the compound name. Names include: L – *cis* hydroxyproline biphenyl ester (*Lc*-BPE); D – *cis* hydroxyproline biphenyl ester (*Dc*-BPE); D – *trans* hydroxyproline biphenyl ester (*Dt*-BPE); L – *trans* hydroxyproline biphenyl ester (*Lt*-BPE).
<sup>b</sup> *Structure* corresponds to the 2D structure of the compound drawn by PerkinElmer ChemDraw
<sup>c</sup> *Docking score* marks the docking score of the top scoring pose using Glide
<sup>f</sup> Corresponds to the value obtained for rASCT2
NA Not experimentally determined

### *Lc*-BPE inhibitory activity with sub-μM potency

The four synthesized compounds were tested for inhibitory activity in rat ASCT2 (rASCT2) with electrophysiology. rASCT2 was used because it exhibits a similar pharmacological profile to that of the human ASCT2, while having better expression in HEK293 cells^15,22,23,29^. In addition, *Lc*-BPE performed similarly against rASCT2 and human ASCT2 (Fig. 2B). Taken together, these results indicate that rASCT2 can serve as a reliable model for ASCT2 inhibitor testing. It is known that competitive inhibitors block a tonic ASCT2 leak anion conductance^29^, which results in the inhibition of inward current in the presence of intracellular permeable anion (SCN^-^, see apparent outward current in Fig. 2A; Extended Data Fig. 2). In contrast, transported substrates activate the anion conductance^30^, resulting in an inward current caused by SCN^-^ outflow (Fig. 2A; Extended Data Fig. 2). The inhibition of the anion leak current was dose dependent and could be fitted to a Michaelis-Menten-type equation to yield an apparent K_i_ of 0.74 ± 0.11 µM for *Lc*-BPE in rASCT2 (Fig. 2B; Extended Data Fig. 2A). In contrast to the L-based diastereomers, the esters based on the D-amino acid scaffold did not show significant inhibition of the leak anion current, indicating that they are not inhibitors of rASCT2 within the concentration range tested (up to 200 μM, Table1). Similar experiments were performed using human ASCT2 with *Lc*-BPE compound yielding a K_i_ of 0.86 ± 0.11 µM (Fig. 2B). Remarkably, the experimental K_i_’s of these newly synthesized compounds correlated with the computational predictions (Table 1). For example, *Lc*-BPE had the best docking score and estimated binding affinity and its docking pose placed the ligand in pocket A, while maintaining the critical interactions with HP1 and TM8, as expected from previously published structures of the human ASCT2^19-21^. Further, the two L-isomers inhibited leak anion current with *Lc*-BPE showing the strongest apparent outward current (Table 1, Extended Data Fig. 2A-C).

**Figure 2.**
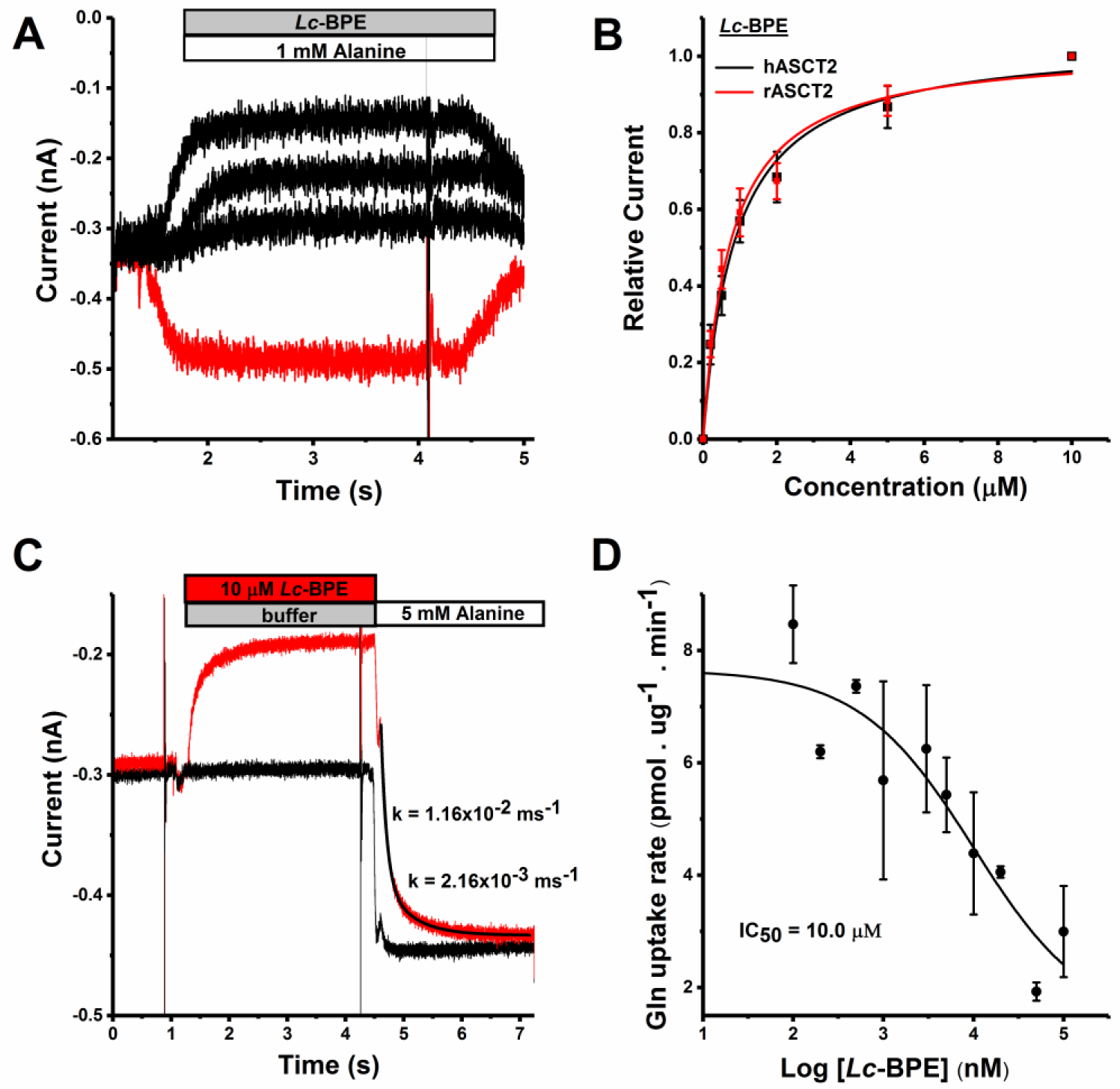
*Lc*-BPE is a potent inhibitor of ASCT2. **(A)** Electrophysiological recordings obtained after application of alanine (red trace) and increasing concentrations of *Lc*-BPE (black traces). The times of application of alanine or *Lc*-BPE are shown by white and grey bars. **(B)** Dose response relationships for the *Lc*-BPE inhibitor. The red (rASCT2) and black (hASCT2) lines represent the best fit to a Michaelis-Menten-like equation with apparent K_i_ of 0.74 µM (rASCT2) and 0.86 µM (hASCT2). Currents were normalized to the current recorded after application of 10 µM inhibitor concentration. **(C)** Current recorded after rapid solution exchange from extracellular buffer (control, gray bar, black trace) to a solution containing buffer + 5 mM alanine (white bar). For the red trace, the cell was pre-incubated with 10 µM *Lc*-BPE (red bar, red trace) followed by rapid application of 5 mM alanine. All current recordings were performed at 0 mV in the presence of 130 mM NaSCN internal and 140 mM NaCl external (homoexchange) solutions. **(D)** Inhibition of glutamine uptake (5 µM external concentration) in the presence of varying concentration of *Lc-*BPE and 5 µM glutamine. IC_50_ of 10.0 µM (black line). Errors bars represent standard deviations from two measurements.

We also tested the specificity of *Lc*-BPE for ASCT2 over other human SLC1 family members (ASCT1, EAAT1-3, and EAAT5). *Lc*-BPE showed inhibitory activity in all SLC1 family members tested, including EAATs, as predicted from the high conservation of pocket A among SLC1 members^22,23,31^. However, the apparent affinity of human ASCT1 (hASCT1) for *Lc*-BPE was approximately 3.8 fold lower than that of rASCT2 (Extended Data Fig. 3A-B), showing slight preference for ASCT2. For the EAATs, the highest affinity interaction of *Lc*-BPE was with hEAAT5 (2.22 ± 0.0.46 µM), or 3-fold higher K_i_ than rASCT2, while interaction with EAATs 1-3 was weaker (Extended Data Fig. 3). This result was not unexpected, because prototypical EAAT inhibitors, such as TBOA^32^, which also have hydrophobic groups in the sidechain that interact with pocket A, are also not entirely selective. Finally, inhibitor activity in a cell proliferation assay with three cancer cell lines, including MCF-7, LnCaP, and MDA-MB-231 was tested. *Lc*-BPE inhibited cell viability in a time- and dose-dependent manner (Extended Data Fig. 4) in the three cell lines, demonstrating its potential pharmacological relevance.

### Inhibitor dissociation kinetics

Due to the potency of *Lc*-BPE, we hypothesized that the inhibitor dissociates slowly from the ASCT2 binding site. To test this, we measured the dissociation kinetics of *Lc*-BPE using a ligand displacement approach. First, rASCT2 expressing cells were pre-incubated with a saturating solution of *L*c-BPE (10 µM) and subsequently the inhibitor was displaced using a super-saturating concentration of the substrate L-alanine (5 mM) (Fig. 2C). It is expected that the inhibitor would have to dissociate before alanine can bind and activate the inward anion current. Consistent with this expectation, activation of the anion current by alanine was slowed significantly after *Lc*-BPE pre-incubation by more than 6-fold. The rate constant for *Lc-*BPE dissociation was estimated as 2.16 s^-1^, indicative of a long residence time of the compound in the rASCT2 binding site.

### *Lc*-BPE is a competitive inhibitor

Our computational models and previous work with proline scaffold compounds predicted *Lc*-BPE as a competitive inhibitor targeting the substrate binding site. To test this prediction, we performed electrophysiological competition experiments in the presence of varying concentrations of the substrate, L-alanine. The results were consistent with a competitive inhibition mechanism. At low inhibitor concentrations, the currents were inward as expected from transported substrates, indicative of the inability of *Lc*-BPE to displace alanine from the binding site. However, as inhibitor concentration increased, the currents became outwardly directed in a dose dependent manner, which marked the displacement of alanine by the inhibitor (Fig. 2C, Extended Data Fig. 5). The apparent K_i_ increased as a function of the alanine concentration, as anticipated for a competitive mechanism (Extended Data Fig. 6,7). We further tested the inhibitory activity of *Lc*-BPE in a cell-free transport assay in proteoliposomes, where reconstituted human ASCT2 catalyzes the exchange of intracellular non-labeled glutamine with extracellular radioactive-labeled glutamine. We observed a dose-dependent inhibition of glutamine transport, when *Lc*-BPE was added externally, which confirms that this compound is an ASCT2 inhibitor with an IC_50_ value of 10 µM in the presence of 5 µM glutamine (Fig. 2D).

### Cryo-EM structure of human ASCT2-*Lc*-BPE

We determined a structure of human ASCT2 reconstituted into lipid nanodiscs, in the presence of the inhibitor *Lc*-BPE, at 3.4 Å resolution using single particle cryo-EM (Fig. 3, Extended Data Fig. 8, Table 2). The protein adopts a single symmetrical state, with all three protomers in the same outward-facing conformation, in which *Lc*-BPE precludes HP2 from closing. In comparison to other ASCT2 structures in outward conformations^21^, the structure showed a distinct HP2 loop conformation that is hinged further away from the binding site by 3 Å compared to the substrate free outward state as measured using the A433 Cα atom (Fig. 3B). *Lc*-BPE overlaps with the region that HP2 occupies in the substrate-bound occluded and substrate-free states, partially displacing HP2. The displacement of HP2 appears to be a consequence of inhibitor binding, suggesting that the HP2 gate is flexible and can potentially move even further away to accommodate bulkier inhibitors. In further support of this hypothesis, the HP2 in this structure, as well as in other ASCT2 structures, is less resolved, indicative of structural flexibility^19-21^. This observation offers a strategy to improve inhibitor potency by designing larger inhibitors to specifically block the gating mechanism of the transporter. In addition, we observed a strong density in the binding site, which overlaps with the density of glutamine in the previous ASCT2 structures^20,21^. This density protruded outside the region occupied by glutamine in the previous structures (Fig. 3C), where *Lc*-BPE is located in a space between HP1 and TM2, revealing a new sub-pocket in the substrate binding site, called pocket C. We refer to this binding mode as a “ligand up” orientation (Fig. 4C and 4D).

**Figure 3.**
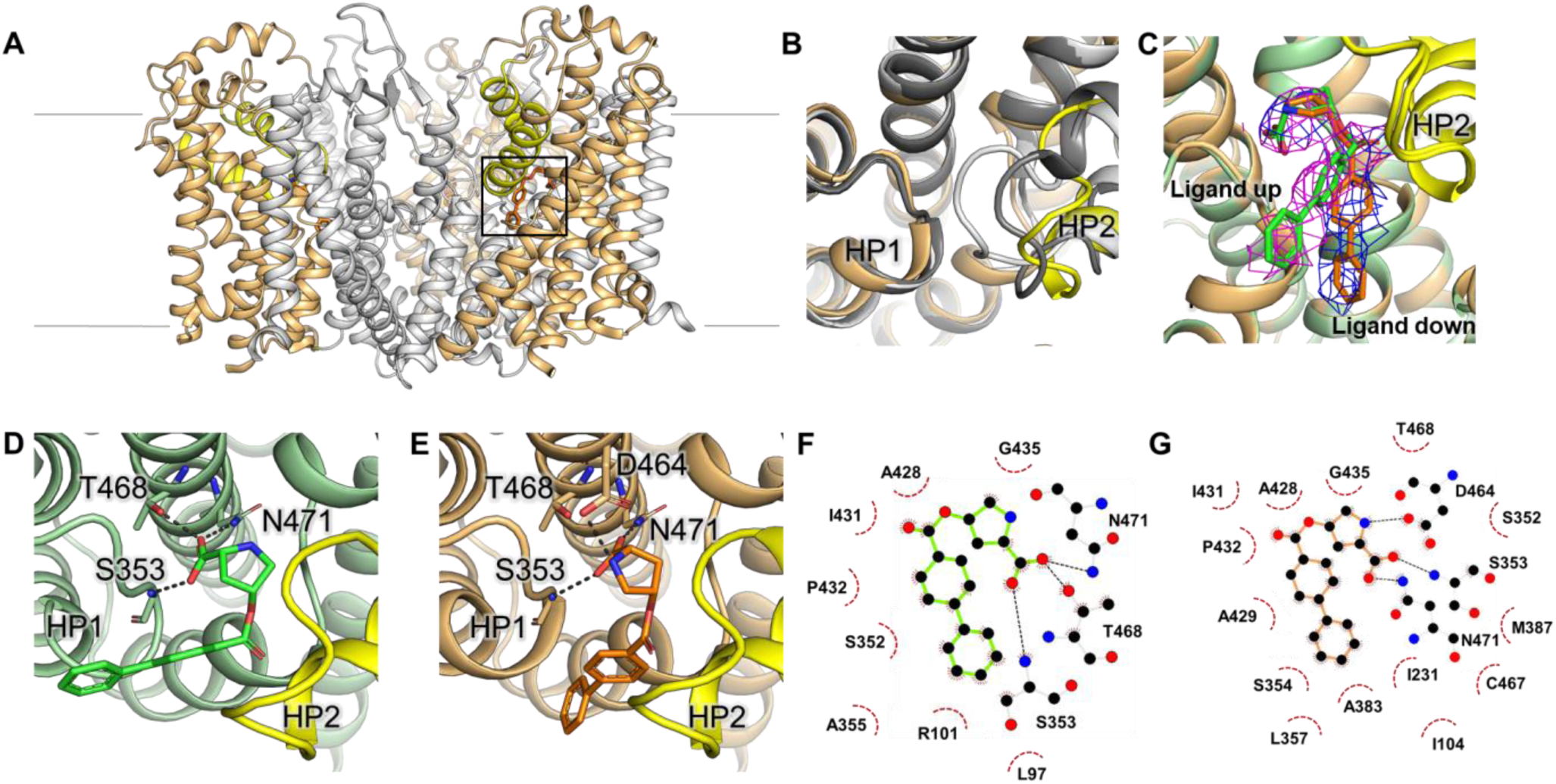
Structural basis for ASCT2 inhibition by *Lc*-BPE. **(A)** Cryo-EM structure of the outward-open ASCT2 trimer in complex with *Lc*-BPE. The scaffold and transport domains are depicted in gray and orange ribbons respectively, with HP2 shown in yellow. Lines show the approximate location of the membrane boundaries. The binding site is roughly outlined with a box. **(B)** Superposition of the apo (6MPB; light gray), outward-occluded (6MP6; dark gray), and outward-open structure in complex with *Lc*-BPE (yellow) of ASCT2, highlighting the HP2 position for the different conformations. **(C)** Zoomed view of the substrate binding site of ASCT2 showing two possible modes of *Lc*-BPE binding. Green structure and ligand with pink density mesh represent the “ligand up” conformation and the orange structure and ligand with blue density mesh represent the “ligand down” conformation. **(D and E)** Potential hydrogen bonds between inhibitor and protein for each ligand conformation shown as dashed lines. In the “ligand up” conformation **(D)** the distal phenyl ring of the ligand interacts with previously unknown sub-pocket. **(F and G)** 2-D ligand interaction plot visualized with LigPlot+^76^ of (**F**) “ligand up” and **(G)** “ligand down” conformations. Hydrogen bonds are represented as black dashes and residues making hydrophobic interactions are marked with red dashes and labeled.

**Figure 4.**
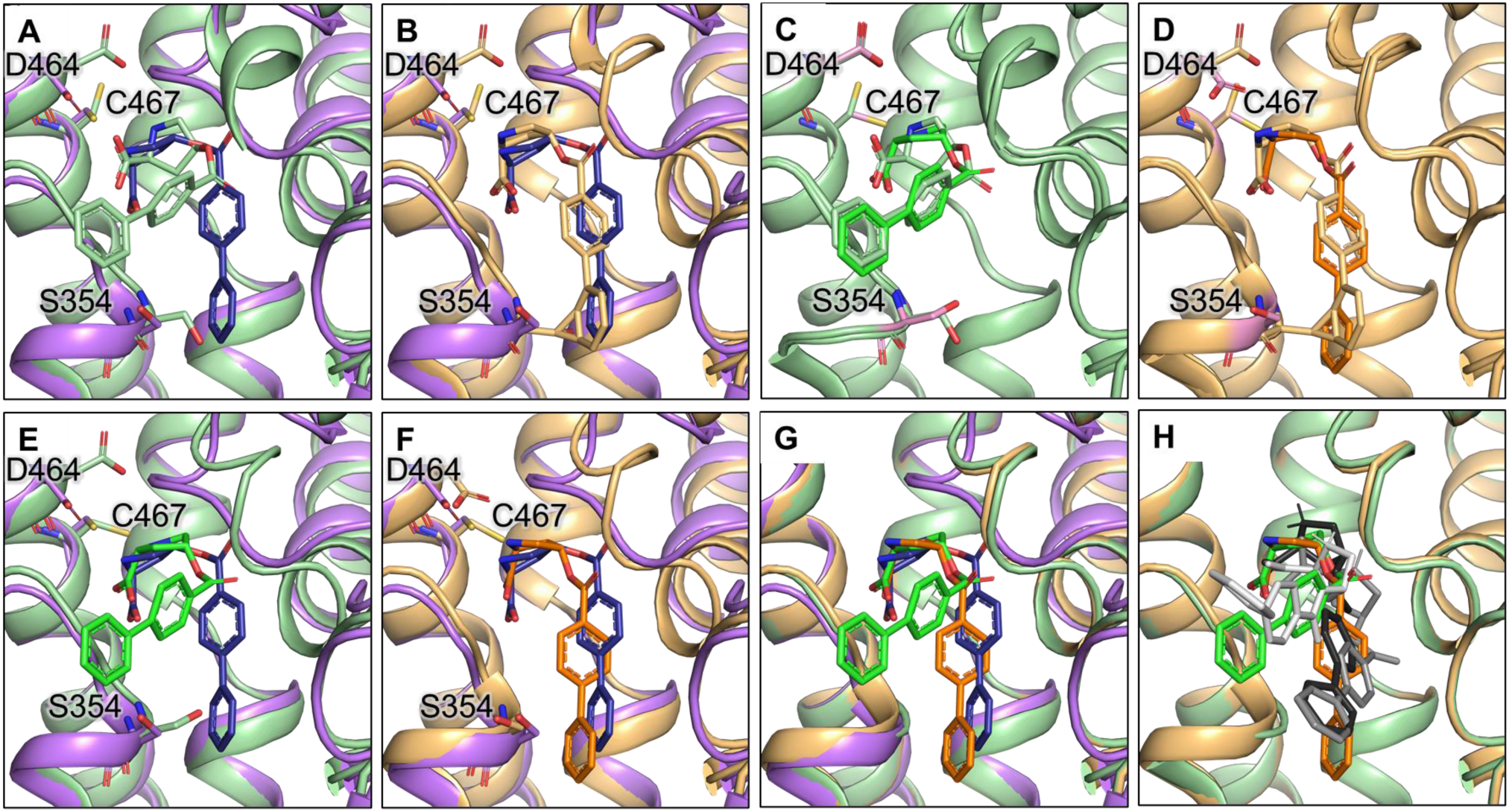
Progressive model building and refinement of the initial fit into the cryo-EM density. Panels show the superposition of the homology model (purple) and cryo-EM structures with “ligand up” (green) and “ligand down” (orange) conformations and highlight the sidechains of S354, D464 and C467 that were remodeled. **(A** and **B)** Superposition of the homology model and initial cryo-EM structures with **(A)** “ligand up” and **(B)** “ligand down” conformations; showing the steric clash between the S354 sidechain and *Lc*-BPE. **(C** and **D)** Superposition of initial and refined structures with remodeled residues in pink sticks. **(C)** “ligand up” initial and refined structures, with the bright green ligand representing the refined *Lc-*BPE position. **(D)** “ligand down” initial and refined structures, with the bright orange representing the refined *Lc-*BPE position. **(E** and **F)** Homology model with docked ligand superposed to **(E)** refined “ligand up” structure and **(F)** refined “ligand down” structure. **(G)** Superposition of refined “ligand up” and “down” structures with the homology model. **(H)** Selected metainference ensemble clusters from by MD simulations with the cryo-EM density as restraint, superposed with the refined “ligand up” and “ligand down” structures. The “ligand up” cluster is light gray and “ligand down” clusters are medium and dark gray, respectively.

**Table 2:**
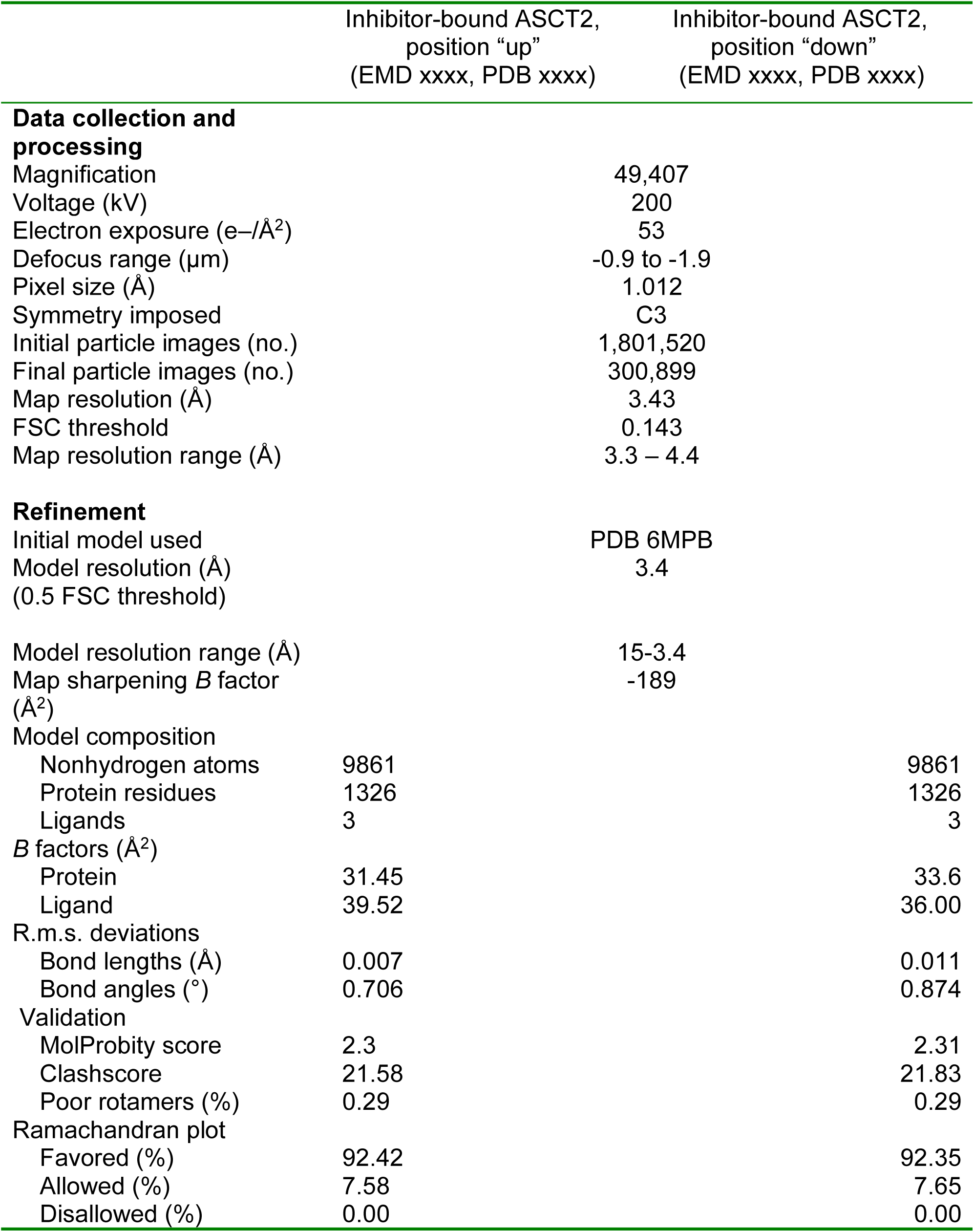
Cryo-EM data collection, refinement, and validation statistics.

Interestingly, the empirically determined “ligand up” orientation was different from the docking pose of *Lc*-BPE in the ASCT2 model (“ligand down”; Extended Data Fig. 1B). Surprisingly, molecular docking using both Glide and OpenEye FRED of *Lc*-BPE against the cryo-EM structure, from which *Lc*-BPE coordinates had been removed, did not recapitulate this “ligand up” binding mode. Upon closer inspection, we identified additional differences between the homology model and the structure initially fitted into the cryo-EM map: in the homology model, S354, D464, and C467 directly interact with the ligand, in agreement with previous observations^18,22,33^. Conversely, in the cryo-EM structure, the sidechain of S354 points in towards the binding site (Fig. 4A), possibly resulting in a steric clash with the ligand and reducing the space in the pocket A available for ligand binding. Further, the sidechain of D464 in the cryo-EM structure pointed up and away from the binding site, thereby reducing the number of critical polar contacts between the ligand and ASCT2 (Fig. 4A).

Based on this analysis, a subsequent exploration of the cryo-EM density map and other nearby unassigned densities, confirmed the possibility of structural rearrangements and an additional “ligand down” conformation for the inhibitor (Fig. 4B). However, because assigning the “ligand down” pose resulted in steric clashes with S354 sidechain, we further compared the cryo-EM structure binding site to structures of other human and prokaryotic and human SLC1 structures in various conformations^10,19-21,34^. This resulted in remodeling the sidechains of S354, C467 and D464 in conformations that resembled those conformations of the corresponding residues in the homology model. Indeed, the refined cryo-EM structure obtained an improved enrichment score (AUC of 79.9 and logAUC 40.4, compared to AUC 69.5 and logAUC 32.4), supporting the remodeling (Fig. 4C-F, Extended Data Fig. 9). Overall, these results indicate the existence of two inhibitor binding modes, “ligand up” and “ligand down”.

Concurrent with previous structures of ASCT2 and other homolog structures with bound ligand, we observe conserved critical polar contacts with the side chain of N471 and the backbone of S353 in HP1 in both “ligand up” and “ligand down” conformations (Fig. 3D-G). In general, “ligand up” *Lc*-BPE is coordinated by S353, T468, and N471 residues, while “ligand down” *Lc*-BPE is coordinated by S353, D464 and N471 residues in the binding site (Fig. 3D-G). Further analysis of the two potential binding modes showed that the “ligand down” state has more favorable hydrophobic interaction at the base of pocket A compared to the “ligand up” state (Fig. 3F and 3G).

### MD simulations confirm multiple binding modes

Areas of lower resolution in cryo-EM maps can be the result of averaging multiple different conformations during image processing. To explore the possibility of multiple pharmacological binding states of inhibitors to ASCT2 we used metainference MD simulations^35^, to model an ensemble of conformations consistent with the cryo-EM data^36^. In metainference, the molecular mechanics force field used in standard MD is augmented by spatial restraints that enforce the agreement of an ensemble of replicas of the system with the cryo-EM density. Using a Bayesian inference framework, this approach accounts for the simultaneous presence of structural heterogeneity, data ensemble-averaging, and variable level of noise in different areas of the experimental map. Metainference has recently been successfully used to characterize the structural heterogeneity of the N-terminal gating region of the ClpP protease^37^ and the effect of acetylation on the dynamics of the K40 loop in α-tubulin^38^.

The ensemble of models obtained with metainference was classified based on the positions of the ligand atoms and those of the surrounding protein residues (Materials and Methods). This analysis indeed suggests that the observed cryo-EM map is the result of an equilibrium of multiple conformations in which the ligand occupies distinct poses and flexible HP2 loop conformations, as anticipated. We observed 6 clusters in total, which we further grouped into 3 distinct groups based on visual analysis (Extended Data Fig 10). These groups support the presence of multiple pharmacologically relevant conformations. Interestingly, the two most populated groups are highly similar to the “ligand up” and “ligand down” configurations, supporting the analysis based on the initial cryo-EM structure and homology model, with the third group being unlikely to be replicated under physiological conditions (Fig. 4H, Extended Data Fig 10). This analysis also supports that HP2 is highly flexible and its position is dependent on the bound inhibitor. We quantified the agreement of the cryo-EM structure and the metainference ensemble with the observed cryo-EM density using the cross-correlation between the experimental and simulated maps. In the former case, the simulated map was calculated on the individual cryo-EM model; in the latter case, the simulated map was averaged over all members of the metainference ensemble. The cross-correlation was computed on the entire experimental grid (CC_box) and was equal to 0.403 and 0.684 in the case of the individual cryo-EM structure and of the metainference ensemble, respectively. Overall, the metainference ensemble provided a better fit of the density map compared to the cryo-EM structure prior to B-factor refinement, without compromising the stereochemical quality of the models (Extended Data Fig. 11).

## DISCUSSION

In this study, we combined computational modeling and experimental testing, to describe the molecular basis for ASCT2 inhibition. Three key findings emerge from our work. First, we developed four ASCT2 inhibitors that provide novel chemical tools for further studying the role of ASCT2 in cancer and other diseases. They include *Lc*-BPE, a competitive, stereoselective inhibitor with sub-μM potency. *Lc*-BPE was shown to bind the substrate binding site of ASCT2 (Fig. 2, Table1). Second, our computational models correctly predicted protein-inhibitor interactions that were validated experimentally by cryo-EM studies and guided structure refinement of a transporter-inhibitor complex (Figs. 3,4). This finding suggests that a similar approach could be applied to solve the structures of other challenging human SLC targets in different conformations. Third, our computational analysis, including iterative model refinement and MD simulations using the cryo-EM density as a restraint, revealed distinct pharmacologically relevant conformations of the ligand and ASCT2 binding site (Figs. 3C, 4). This result describes the mechanism by which ASCT2 inhibitors achieve their stereospecificity and provides a framework for developing future clinically active, ASCT2-selective compounds. We discuss each of these three findings in turn.

### Novel chemical tools to characterize ASCT2

A chemical tool is a molecule capable of modulating the function of a protein, enabling researchers to ask mechanistic questions about its molecular target with various experimental systems, including biochemical, cellular, or *in vivo* approaches. Notably, a useful chemical probe needs to be validated experimentally using orthogonal assays^39^. Although the SLC1 family has been established as an important family of druggable proteins, the SLC1 members, including ASCT2, currently suffer from a limited number of chemical probes to help decipher their role in disease. Here we developed four ASCT2 inhibitors that are ester derivatives based on the previously characterized 4-hydroxyproline amino acid scaffold. The isomer with the highest apparent affinity, *Lc*-BPE, was validated using electrophysiological, cellular, biochemical, and structural approaches directly showing target engagement and confirming a competitive stereoselective inhibition mechanism. In addition, as predicted from our models, *Lc*-BPE, which interacts with a conserved pocket across all human SLC1 members (pocket A), was not found to be strongly selective for ASCT2 over other SLC1 members (Extended Data Fig. 3). While this will likely limit the clinical relevance of *Lc*-BPE, the compound provides an excellent starting point for designing ASCT2-specifc compounds, such as those targeting specific subpockets of this protein (Fig. 3C and Extended Data Fig. 1A). With these limitations in mind, it should be noted that, to our knowledge, *Lc*-BPE is the first inhibitor for ASCTs with a sub-μM affinity. Thus, this compound represents a major step towards inhibitors with high affinity ASCT2 interaction.

### Accurate prediction of transporter-inhibitor interaction for a challenging target

Despite progress in methodologies for structure determination of membrane proteins, the coverage of human SLC structures is limited. Particularly, structures of only a limited number human SLC transporters have been determined at high resolution^31^. Furthermore, while the SLCs are highly dynamic, most proteins with known structures have not been characterized in multiple conformations. Therefore, homology modeling can be an efficient and accurate method to visualize SLC transporters in different conformations. Recently, the combination of different sampling and refinement approaches with ligand docking has aided the design of small molecule ligands, particularly, for challenging membrane protein targets^19,31^. Here iterative modeling and molecular docking has enabled us to accurately model the ASCT2 structure. The final homology model, based on the human EAAT1 structure, obtained high ASCT2 ligand enrichment and exhibited high structural similarity to the later experimentally solved cryo-EM structure. The homology model also guided precise cryo-EM model refinement, highlighting not only its accurate prediction power but also its versatility as a complementary approach to empirical structure determination. During refinement, we focused on three positions that coordinate *Lc*-BPE binding, S354, D464, and C467, which have been shown to be critical for ASCT2 substrate specificity^19-21^. Overall, we demonstrate a quick and efficient approach to use homology models in a previously uncharacterized conformation, to explore novel chemical spaces in drug design. Notably, this approach is generally applicable to characterize other challenging membrane protein targets.

### Integrated approach identifies multiple pharmacologically relevant states

Cryo-EM has become the most dominant technology for structure determination of human membrane proteins, particularly SLC transporters^40^. Computational methods such as various MD simulation methodologies can efficiently sample relevant conformations and expedite structure determination with cryo-EM^41,42^. In this work, the integration of multiple computational methods with varying levels of speed and robustness throughout the structure determination process aided the characterization of ASCT2-ligand interactions. First, homology models of ASCT2 in the outward-facing conformation enabled the rational design of compounds targeting pocket A of the substrate binding site. The newly developed compounds have aided structure determination of this outward-facing state of ASCT2 in which HP2 is in an open conformation (e.g., Fig. 3B).

Second, computational modeling during the cryo-EM structure determination process, has progressively improved the quality of the solved structure. In particular, combining molecular docking and ligand enrichment informed cryo-EM model refinement, ultimately leading to the identification of an additional pharmacologically relevant state. The initial structure revealed a conformation that fits the cryo-EM density map with *Lc*-BPE pointing into the space between HP1 and TM2 (“ligand up”), which previously was not thought to comprise the ASCT2 substrate binding site (Fig. 3C). Further analysis of the density map identified the “ligand down” conformation that strikingly resembled the docking pose in the homology model. This result suggests that the model represents a pharmacologically relevant conformation, as was previously suggested for GPCRs using other approaches^43^. Third, using MD simulations with the density map as a restraint, revealed multiple potential ASCT2-*Lc*-BPE configurations, which can be further grouped into 3 putative positions of the ligand (Fig. 4H and Extended Data Fig. 10). Interestingly, two of these three positions correspond to conformations similar to the ‘ligand up’ and ‘ligand down’ arrangements indicated from the electron densities, increasing our confidence in this approach. Notably, these two configurations could guide the exploration of distinct chemical spaces around *Lc*-BPE in order to improve its affinity and selectivity for the development of ASCT2-targeted drugs.

In summary, this study described the mechanism of transport inhibition in a pharmacologically relevant transporter target, ASCT2, by combining modern methods in protein structure determination and computational modeling. The results provide novel chemical tools to further characterize the role of ASCT2 in disease, and a framework for designing future clinically relevant compounds. The approach presented in this work is generally applicable to other challenging membrane protein targets such as the human SLC transporters.

## MATERIALS AND METHODS

### ASCT2 modeling and assessment

We have previously developed homology models of ASCT2 based on the outward-open structures of the human EAAT1 transporter^25,27^. Here, we iteratively refined this model based on its ability to discriminate ligand from decoy compounds with ligand enrichment calculations. The sidechains of the binding site residues S354, D464, and C467 were remodeled on fixed backbone using PyMOL^44^ and SCRWL4^45^, guided by recently determined structures of ASCT2 in multiple conformations. PyMOL mutagenesis was done with backbone dependent rotamers and SCRWL4 was done according to default parameters. Enrichment was done using a library of 29 known ASCT2 ligands, including substrates and inhibitors that were collected from the literature^15,16,24,46-48^ and ChEMBL^49^ and 1,434 decoys generated with the DUD-E server^50^. Docking was performed with OpenEye FRED^51^ as described in our previous work^27^. Hydrogen-bond constraints used for docking against the model at HP1; S351 and S353, and the refined structure at HP1; S353 and TM8; N471.

### Molecular docking with Schrödinger

Docking of the four ligands was performed with the Schrödinger suite using Glide^52^. In brief, the models were prepared for docking using the Maestro Protein Preparation Wizard under default parameters. The substrate binding site was defined using the Maestro Receptor Grid Generation panel and the coordinates of the reference ligand (TFB-TBOA) were derived from the EAAT1 template structure. The allosteric ligand UCPH_101_ was removed from the model prior to defining the site, to avoid multiple reference ligands in the grid. Docking was conducted without constraints. Compounds were prepared for docking with Glide v19-3 with LigPrep under default parameters^52^.

### Relative binding affinity prediction

We estimated the relative binding affinity between the compounds and the ASCT2 model binding site with mechanics generalized with born surface area solvation (MM-GBSA). These calculations were relative, where a more negative value was indicative of a better binding affinity. Here we used MM-GBSA with Prime from the Schrödinger suite v18-4)^52^. The model was prepared as described above with the exception that reference ligands were removed prior, and we used the docked ligands from above as input. Standard parameters were used, with the distance from the ligand set to 5 Å for all calculations.

### MD simulations

The atoms of chain A and of the associated ligand were extracted from the ‘ligand up’ cryo-EM structure. CHIMERA^53^ was used to select the cryo-EM density within 5 Å of the model to be used as input of the metainference simulation of the ASCT2 monomer. The starting model was prepared using CHARMM-GUI^54^. 113 POPC lipids were added to the system along with 11554 water molecules in a triclinic periodic box of volume equal to 616 nm^3^. 31 potassium and 30 chloride ions were added to ensure charge neutrality and a salt concentration of 0.15 M. The CHARMM36 force field^55^ was used for the protein, lipids, and ions; the CHARMM General Force Field and the TIP3P model were used for the ligand and water molecules, respectively. A 30 ns-long equilibration was performed following the standard CHARMM-GUI protocol consisting in multiple consecutive simulations in the NVT and NPT ensembles. During these equilibration steps, harmonic restraints on the positions of the lipids, ligand, and protein atoms were gradually switched off.

In the metainference simulation, a Gaussian noise model with one error parameter for each voxel of the cryo-EM map was used. These variables were marginalized to avoid their explicit sampling, as done in previous applications^36,37^. 16 replicas of the system were used, and their initial configurations were randomly selected from the last 10ns-long step of the equilibration protocol. The metainference run was conducted for a total aggregated time of 1.8 μs. All simulations were carried out using GROMACS 2019.6^56^ and the ISDB module^41^ of the open-source, community-developed PLUMED^57^ library (GitHub ISDB branch; https://github.com/plumed/plumed2/tree/isdb). For the analysis, the initial frames of the trajectory of each replica, corresponding to 20% of the total simulation time, were considered as additional equilibration steps under the cryo-EM restraint and discarded. The remaining conformations from all replicas were merged together and clustered using: i) the Root Mean Square Deviation calculated on all the heavy atoms of the ligand and of the protein residues within 5 Å of the ligand in at least one member of the ensemble; ii) the gromos algorithm^58^ with a cutoff equal to 2.5 Å. PDB of the starting model, topology files, GROMACS inputs for equilibration and production runs, PLUMED input files for the metainference simulation, and analysis scripts are available on PLUMED-NEST^57^ (www.plumed-nest.org) under accession code plumID:20.015.

### ASCT2 expression and purification

Human ASCT2 (hASCT2) was produced in *Pichia pastoris* X-33 strain (Invitrogen) in fermentor^59^ and purified using DDM and CHS (Anatrace) and 1 mM L-glutamine (Merck) to maintain protein stability^20^. Membranes representing ∼1.5 g cells were solubilized in buffer A (25 mM Tris-HCl, pH 7.4, 300 mM NaCl, 10% (vol/vol) glycerol, 1 mM L-glutamine, 1% DDM and 0.1% CHS) for 1 h at 4 °C; ultracentrifuged (30 min, 442,907 × g, 4 °C) and supernatant containing solubilized protein was incubated with Ni^2+^-Sepharose resin for 1 h at 4 °C. Protein was eluted with buffer B (20 mM Tris-HCl, pH 7.4, 300 mM NaCl, 500 mM imidazole, pH 7.4, 10% glycerol, 1 mM L-glutamine, 0.02% DDM and 0.002% CHS), and applied to size-exclusion chromatography with a Superdex 200 10/300 gel-filtration column (GE Healthcare) preequilibrated with buffer C (20 mM Tris-HCl, pH 7.4, 300 mM NaCl, 1 mM L-glutamine, 0.02% DDM and 0.002% CHS). Peak elution fractions were immediately used in further procedures.

### Reconstitution into proteoliposomes and transport assays

Freshly purified ASCT2 was reconstituted in the liposomes composed of *Escherichia coli* polar lipids and egg phosphatidylcholine at a 3:1 ratio (w/w) and supplemented with 10% (w/w) cholesterol (Avanti Polar Lipids)^20^. For transport assays proteoliposomes were loaded with 50 mM NaCl and 10 mM or 5 mM glutamine using three freeze-thawing cycles, then extruded 11 times through a 400-nm-diameter polycarbonate filter (Avestin), diluted in buffer D (20 mM Tris pH 7.0) and collected during ultracentrifugation (45 min, 442,907 × g, 4 °C). Proteoliposomes were resuspended in buffer D (∼1 μg protein per 1.5 μl) and used in the transport assays carried out in a water bath at 25 °C with constant stirring. Transport was initiated by dilution of 1.5 μl proteoliposomes in 80 μl external buffer (50 mM NaCl and 50 μM or 5 μM [^3^H]glutamine (PerkinElmer) in 20 mM Tris pH 7.0). Inhibitors or equivalent amounts of DMSO were added to the external buffer mixture. At indicated time points the reaction was stopped by diluting the mixture in 2 ml of cold buffer D, filtered over a 0.45-μm pore-size filter (Portran BA-85, Whatman), washed with 2 ml of cold buffer D and filtered again. The level of radioactivity accumulated inside the proteoliposomes, as a consequence of amino-acid exchange, was counted using a PerkinElmer Tri-Carb 2800RT liquid scintillation counter after dissolving the filter in 2 ml of scintillation liquid (Emulsifier Scintillator Plus, PerkinElmer).

### Reconstitution of ASCT2 in nanodiscs

An aliquot of mixture of *E*.*coli* polar lipids and egg phosphatidylcholine (3:1, w/w) supplemented with 10% (w/w) cholesterol was solubilized with 30 mM DDM-CHS for 3 h while nutating. Nanodiscs were assembled at a reconstitution ratio of 1 nmol ASCT2 (as monomer): 5 nmol of MSP2N2: 70 nmol of solubilized lipids. For this freshly purified ASCT2 was first mixed with solubilized lipids and incubated for 30 min at 4 °C. Purified and TEV protease treated MSP2N2^60^ was added for the following 30 min. 300 mg of SM2 BioBead (Bio-Rad) were added overnight to remove DDM. Assembled nanodiscs were purified from discs devoid of ASCT2 via Ni^2+^-Sepharose chromatography, collected in elution fraction in buffer E (20 mM Tris-HCl, pH 7.4, 300 mM NaCl, 500 mM imidazole, pH 7.4) and further applied on size-exclusion chromatography with a Superdex 200 10/300 gel-filtration column preequilibrated with buffer F (20 mM Tris-HCl, pH 7.4, 200 mM NaCl).

### Cryo-EM sample preparation and data collection

Freshly purified hASCT2 nanodiscs were concentrated to ∼1 mg ml^−1^ using an Amicon Ultra-0.5 mL concentrating device (Merck) with a 100 kDa filter cut-off and then 100 μM inhibitor was added and incubated for 1 h on ice. 2.8 μl of the sample were applied onto the holey-carbon cryo-EM grids (Au R1.2/1.3, 300 mesh, Quantifoil), which were preliminary glow discharged at 5 mA for 30 s, blotted for 3–4 s in a Vitrobot Mark IV (Thermo Fisher) at 15 °C and 100% humidity, plunge frozen into a liquid ethane/propane mixture and stored in liquid nitrogen until further use. Screening of the grid areas with best ice properties was done with the help of a self-written script to calculate the ice thickness. Cryo-EM data in selected grid regions were collected in-house on a 200-keV Talos Arctica microscope (Thermo Fisher) with a post-column energy filter (Gatan) in zero-loss mode, with a 20-eV slit and a 100-μm objective aperture. Images were acquired in an automatic manner with EPU (Thermo Fisher) on a K2 summit detector (Gatan) in counting mode at ×49,407 magnification (1.012 Å pixel size) and a defocus range from −0.9 to −1.9 μm. During an exposure time of 9 s, 60 frames were recorded with a total dose of about 53 electrons/Å^2^. On-the-fly data quality was monitored using FOCUS software^61^.

### Image processing

For the ASCT2 nanodiscs dataset in the presence of inhibitor, 6,233 micrographs were recorded. Beam-induced motion was corrected with MotionCor2_1.2.1^62^ and the CTF parameters estimated with ctffind4.1.13^63^. Recorded micrographs were manually checked in FOCUS (1.1.0), and micrographs, which were out of defocus range (<0.4 and >2 μm), contaminated with ice or aggregates, and with a low-resolution estimation of the CTF fit (>4 Å), were discarded. The remaining 5,991 micrographs were imported in cryoSPARC v2.14.2^64^. Around 1000 particles were manually picked to create templates for particle autopicking. 3,666,842 particles were autopicked and extracted with a box size of 200 pixels. After 2D classification 1,801,520 particles were left and majority of particles (1,427,784) were used for several rounds of ab-initio volume generation, and C3 symmetry was applied. 404,284 particles of best classes and 373,736 particles remaining after 2D classification (778,020 particles in total) were exported from cryoSPARC and imported in RELION-3.0.8^65^ and used in 3D classification with C3 symmetry applied and resulted in a one best class with 300,899 particles (38.7%). These particles were used in the refinement job, where hASCT2 map generated in cryoSPARC was used as a reference and was low-pass filtered to 15 Å, and C3 symmetry was applied. In the last refinement iteration, a mask excluding nanodisc was used and the refinement continued until convergence (focus refinement), following postprocessing job, which resulted in a map at 4 Å. Four rounds of per-particle CTF refinement and beam tilt refinement in Relion3^8^ improved resolution to 3.43 Å.

To check for conformational heterogeneity, we performed 3D classifications without imposed C3 symmetry at different stages of image processing, and we did not find other conformations present. We also did 3D classifications of individual protomers after symmetry expansion and signal subtraction to check for conformational heterogeneity within the trimer. All particles were clustered in one class indicating the presence of only one protein conformation within the trimer. Bayesian polishing in RELION3^66^ did not lead to further improvement in map resolution. The resolution was estimated using the 0.143 cut-off criterion^67^ with gold-standard Fourier shell correlation (FSC) between two independently refined half-maps^68^. During post-processing, the approach of high-resolution noise substitution was used to correct for convolution effects of real-space masking on the FSC curve^69^. The directional resolution anisotropy of density map was quantitatively evaluated using the 3DFSC web interface (https://3dfsc.salk.edu)^70^.

### Cryo-EM model building and validation

Model was built in COOT^71^ using the previously determined ASCT2 structure^21^ in detergent as reference. The resolution of the map was of a good quality to unambiguously assign the protein sequence and model most of the residues (47–489). Blurring of the final map to b-factors −100 Å^2^ and −50 Å^2^ helped to control loops fitting. Real-space refinements were performed in Phenix^72^ with NCS restrains option. The quality of the fit was validated by a Fourier shell cross correlation (FSCsum) between the refined model and the final map. To monitor the effects of potential overfitting, random shifts (up to 0.5 Å) were introduced into the coordinates of the final model, followed by refinement against the first unfiltered half map. The FSC between this shaken-refined model and the first half map used during validation refinement is termed FSCwork, and the FSC against the second half map, which was not used at any point during refinement, is termed FSCfree. A marginal gap between the curves describing FSCwork and FSCfree indicates no overfitting of the model.

The SBGrid software package tool was used^73^. Images were prepared with PyMOL^44^, ChimeraX^73^, Chimera^53^.

### Cell culture and transfection

Human embryonic kidney 293 (HEK293, ATCC CRL-11268) cells were cultured in DMEM media supplemented with 10% (*v/v*) fetal bovine serum (FBS), 2 mM glutamine, 1% penicillin streptomycin solution, 1 mM sodium pyruvate and non-essential amino acids. Cells were maintained at 37°C in a fully humidified atmosphere containing 5% CO_2_. rASCT2, hASCT2, hASCT1, EAAT1, EAAT2, EAAT3, EAAT5 and YFP complementary DNAs were each used to transiently transfect HEK293 using POLYPLUS Jet-prime transfection reagent. Cells were analyzed using electrophysiological techniques 24-40 hours after transfection.

Human breast cancer cell lines MDA-MB-231 (kindly provided by Dr. Tracy Brooks, Binghamton University) and human prostate cancer cell line LnCaP (ATCC. CRL-1740) were cultured in RPMI-1640 media supplemented with 10% (*v/v*) FBS, 2 mM glutamine, 1% (*v/v*) penicillin streptomycin solution and 1mM sodium pyruvate. MCF-7 cells (kindly provided by Dr. Tracy Brooks, Binghamton University) were cultured in MEM media supplemented with 10% (*v/v*) FBS, 2 mM glutamine, 1% (*v/v*) penicillin streptomycin solution, 1 mM sodium pyruvate and non-essential amino acids enriched with 0.01 mg/mL insulin.

### Cell viability assays

2 × 10^3^ to 10^4^ cells per well were plated in flat-bottomed 96-well plates and allowed to adhere for 12 hours before treatment. Cells were treated with or without benzyl serine or inhibitors and incubated for up to 72 hours. Proliferation was measured at days 0, 1, 2, and 3 using the MTT (3-(4,5-dimethylthiazol-2-yl)-2,5-diphenyltetrazolium bromide) assay. In brief, 10 μL MTT solution (5 mg/mL) was added to each well and plates incubated for 5-6 h. MTT solution was removed and 100 μL DMSO added to each well and swirled gently and incubated for 30 minutes. Absorbance in each well was read at both 570 nm and 630 nm using Synergy HTX plate reader (BioTek Instruments). Results were then plotted as percentages of the absorbance of control cells. Statistical significance of data was measured using a paired sample t-test where, p<0.05 = *, p<0.01 = **, p<0.001 = *** and p<0.0001 = ****.

### Electrophysiological techniques

Electrophysiological experiments were performed as described previously^15,29^. Stock solutions of inhibitor was prepared in dimethyl sulfoxide (DMSO) up to 100 mM. Dilutions to working concentrations were made using external buffer. The highest DMSO concentration used (2%) did not affect electrophysiological results, as shown in control cells (data not shown). For rASCT2, hASCT2 and hASCT1, external buffer contained 140 mM NaCl, 2 mM MgCl_2_, 2 mM CaCl_2_, and 10 mM HEPES, pH 7.40 while internal pipette solution comprised of 130 mM NaSCN, 2 mM MgCl_2_, 10 mM EGTA, 10 mM HEPES and 10mM alanine, pH 7.40. For specificity experiments with EAAT1, EAAT2, EAAC1 and EAAT5, internal solution contained 10 mM glutamate instead of alanine. Compounds were applied to HEK293 cells expressing DNA of interest suspended from a current recording electrode in whole cell configuration^74^ through a rapid solution exchange device described previously^75^. Cells are immersed in external buffer bath used to dissolve the inhibitors. The open pipette resistance was between 3 and 6 MΩ. Series resistance was not compensated in these experiments due to relatively small currents. Currents traces were recorded using an Adams and List EPC7 amplifier and digitized using a Molecular Devices Digidata A/D converter.

### Data analysis

Data analysis was performed as described previously^23^. Linear and nonlinear curve fitting of the experimental data were analyzed using MicroCal Origin software. Linear plots were fitted using the general equation (y = a + bx) obtaining adjusted R^2^ and Pearson’s r values. Nonlinear dose–response relationships were fitted with a Michaelis–Menten-like equation to obtain apparent K_i_ values in the absence of substrate. For competition studies, equation *I* = *I*_1_ + *I*_2_ [Inh]/(K_i_ + [Inh]) was used for fitting, where *I*_1_ is the alanine induced current without the inhibitor, and *I*_2_ is the maximum current in the presence of saturating inhibitor concentration, [Inh]_max_^15,29^. At least four experiments were performed with at least three different cells. Unless stated otherwise, the error bars in all our graphs represent mean ± SD.

### Synthesis

Chemicals were purchased from VWR or Sigma-Aldrich. Except for one isomer, all other compounds were synthesized following the same general procedure as shown in the reaction scheme.

**Scheme 1:**
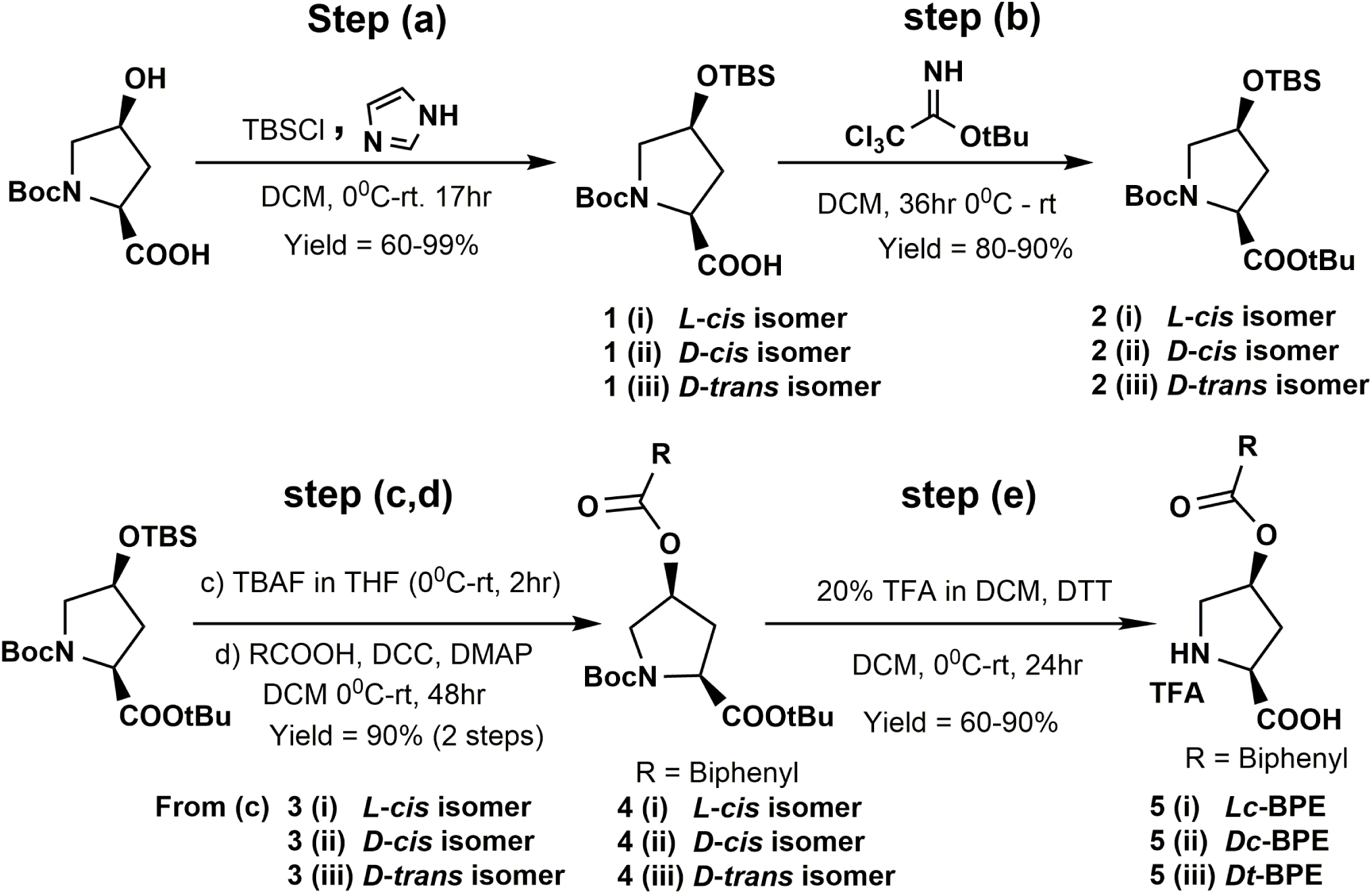
General synthesis of hydroxyproline biphenyl ester compounds. (a) TBSCl, imidazole, DCM, 17h, yield 99%. (b) tert-Butyl 2,2,2-trichloroacetimidate, DCM, 36 h, yield 80-90%. (c) TBAF, THF, 2h, (d) biphenyl carboxylic acid, DCC, DMAP, DCM, 48h, total yield 90%. (e) DTT, 20% TFA in DCM, yield 60%.

**Scheme 2:**
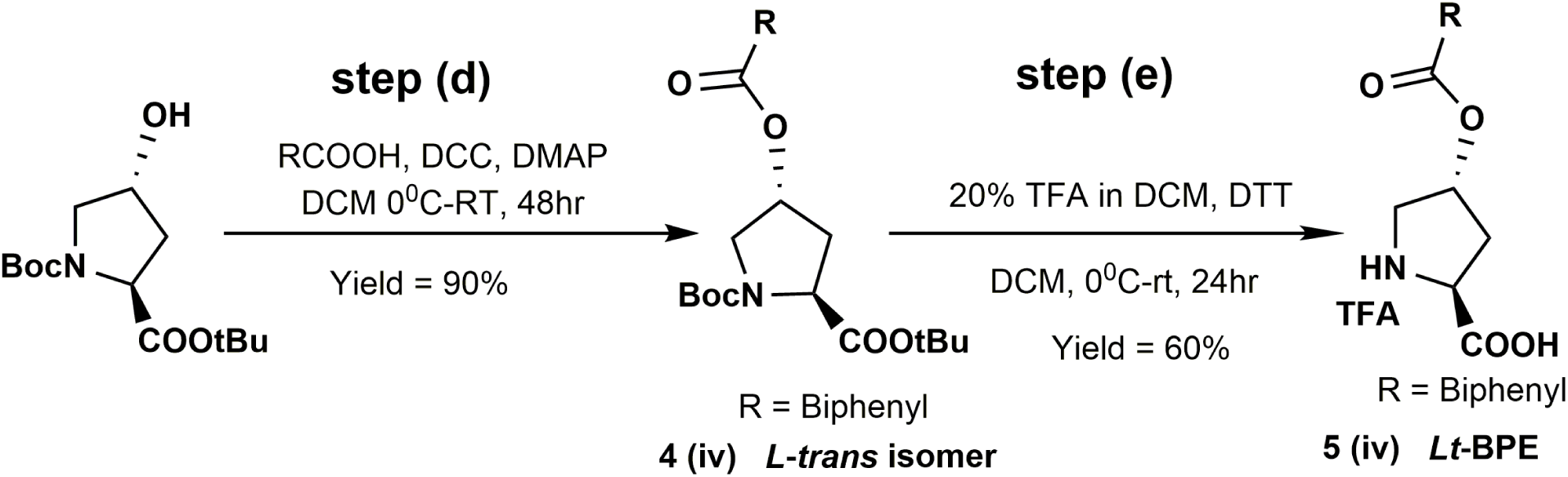
Synthesis of L – *trans* hydroxyproline biphenyl ester compound. (d) 4-biphenyl carboxylic acid, DCC, DMAP, DCM, 48h, total yield 90%. (e) DTT, 20% TFA in DCM, yield 60%.

### General synthesis procedures

**Step (a):** (tert-butoxycarbonyl)-4-hydroxypyrrolidine-2-carboxylic acid (500 mg, 2.16 mmol) and imidazole (744 mg, 10.80 mmol) were weighed into an oven-dried round bottomed flask (RBF) and dissolved in dry DMF (6 mL). The reaction mixture was cooled to 0 °C and TBSCl in dry DMF (652 mg, 4.32 mmol) was added dropwise under N_2_ gas. The reaction mixture was then left to warm up to room temperature and stirred for 24 hours. After removal of excess DMF using N_2_ gas at 50 °C, the residue was suspended in ethyl acetate and washed twice with water, thrice with chilled 1 M HCl and once with brine. The organic layer was dried over anhydrous Na_2_SO_4_ and filtered. The filtrate was concentrated under reduced pressure to a colorless oil. The oil was dissolved in methanol (3 mL) and THF (4 mL) and the solution cooled to 0 °C. LiOH.H_2_O (227 mg, 5.40 mmol) in water (3 mL) was added dropwise and the mixture was allowed to warm to room temperature and stirred for 2 hours. The pH of the solution was adjusted to 2-3 using chilled 1 M HCl and the product was collected as a pure, white precipitate after suction filtration.

**Step (b):** (*tert*-butoxycarbonyl)-4-((*tert*-butyldimethylsilyl)oxy)pyrrolidine-2-carboxylic acid (661 mg, 1.91 mmol) was dissolved in dry DCM and cooled to 0 °C. 1.03 mL of *tert*-Butyl 2,2,2-trichloroacetimidate (1,252 mg, 5.73 mmol) was added dropwise under N_2_ and the mixture was stirred for 36 hours at room temperature. Excess solvent was removed *in vacuo* and the residue purified using flash silica gel chromatography (0 – 15% ethyl acetate in hexanes) to obtain a pure colorless oil.

**Step (c):** di-*tert*-butyl 4-((*tert*-butyldimethylsilyl)oxy)pyrrolidine-1,2-dicarboxylate (635 mg, 1.58 mmol) was dissolved in dry THF and cooled to 0 °C. 1M TBAF in THF (2.05 mL, 2.05 mmol) was added dropwise under N_2_ atmosphere and reaction left to stir for 2 hours. Contents were diluted 25% ethyl acetate in hexanes and washed saturated NH_4_Cl (1x), chilled 0.5M HCl (1x), NaHCO_3_ (1x) and 1:1 mixture of H_2_O and brine (1x). The organic layer was dried over Na_2_SO_4_ and filtered. The filtrate was concentrated *in vacuo* and the colorless oil formed was used in the next step without further purification. For **3 (i)** L – *cis* isomer, the product was purified using flash silica gel chromatography (25 – 60% ethyl acetate in hexanes) to obtain a colorless oil which was precipitated from DCM with hexanes to obtain a pure white solid.

**Step (d):** di-*tert*-butyl 4-hydroxypyrrolidine-1,2-dicarboxylate (52 mg, 0.18 mmol), DMAP (4 mg, 0.04 mmol), [1,1’-biphenyl]-4-carboxylic acid (144 mg, 0.72 mmol) were dissolved in dry DCM and cooled to 0 °C. DCC in dry DCM (75, 0.36 mmol) was added dropwise under N_2_ gas and the reaction mixture was allowed to warm up to room temperature and was stirred for 48 hours. The contents were filtered off and the filtrate concentrated *in vacuo*. The residue was suspended in 50% ethyl acetate in hexanes and washed with NaHCO_3_ (3x), chilled 0.5M HCl (3x), brine (3x) and H_2_O (1x). The organic phase was collected and dried over Na_2_SO_4_ and filtered. The filtrate was concentrated *in vacuo* and the product purified using flash silica gel chromatography (10 – 50% ethyl acetate in hexanes) to obtain a pure white solid.

**Step (e):** di-*tert*-butyl 4-(([1,1’-biphenyl]-4-carbonyl)oxy)pyrrolidine-1,2-dicarboxylate (74 mg, 0.16 mmol) and DTT (49 mg, 0.32 mmol) were weighed into a 10 mL long-necked RBF and dissolved in dry DCM. Trifluoroacetic acid (TFA) (0.39 mL, 5.12 mmol) was added dropwise under N_2_ gas and the reaction mixture was stirred at room temperature for 36 hours. TFA was completely removed under reduced pressure and the product was collected as a white solid after trituration in methanol and diethyl ether. The product was confirmed by TLC (20 - 40% methanol in DCM), rf ∼ 0.2) and was used in all experiments without further purification.

### NMR data for major intermediates

*(2S,4S)-1-(tert-butoxycarbonyl)-4-((tert-butyldimethylsilyl)oxy)pyrrolidine-2-carboxylic acid* **1 (i)**. Synthesized according to general procedure for step (a). Yield 70%, white solid. ^1^H NMR (Chloroform-*d*, 400 MHz) δ 11.06 (1H, s), 4.52 – 4.10 (2H, m), 3.58 (1H, d, *J*=14.5 Hz), 3.47 – 3.10 (1H, m), 2.20 (2H, dd, *J*=61.1, 26.1 Hz), 1.42 (9H, s), 0.84 (9H, s), 0.05 (6H, s). ^13^C NMR (Chloroform-*d*, 101 MHz) δ 176.91, 154.12, 80.77, 70.21, 58.06, 54.48, 39.26, 28.41, 25.72, 18.02, −4.87.

*(2R,4R)-1-(tert-butoxycarbonyl)-4-((tert-butyldimethylsilyl)oxy)pyrrolidine-2-carboxylic acid* **(1 (ii)**. Prepared following general procedure as in **1 (i)** above. Yield 60%, white solid. ^1^H NMR (Chloroform-*d*, 400 MHz) δ 10.24 (1H, s), 4.39 (2H, d, *J*=22.7 Hz), 3.68 – 3.18 (2H, m), 2.21 (2H, s), 1.48 (9H, s), 0.89 (9H, s), 0.11 (6H, s). Full ^13^C NMR could not be obtained due to poor resolution of carbonyl carbons for major experiments done such as routine ^13^C NMR and HMBC.

*(2R,4S)-1-(tert-butoxycarbonyl)-4-((tert-butyldimethylsilyl)oxy)pyrrolidine-2-carboxylic acid* **1 (iii)**. Compound was synthesized following the general procedure for step (a). Yield 99%, white solid. ^1^H NMR (Chloroform-*d*, 400 MHz) δ 9.71 (1H, s), 4.56 – 4.24 (2H, m), 3.71 – 3.27 (2H, m), 2.41 – 2.15 (1H, m), 2.09 (1H, tt, *J*=8.5, 4.5 Hz), 1.44 (9H, d, *J*=20.8 Hz), 0.87 (9H, s), 0.06 (6H, d, *J*=2.2 Hz). ^13^C NMR (Chloroform-*d*, 101 MHz) δ 175.07, 157.13, 81.78, 70.13, 58.66, 55.11, 37.79, 28.50, 25.83, 18.07, −4.70.

*(2S,4S)-di-tert-butyl 4-((tert-butyldimethylsilyl)oxy)pyrrolidine-1,2-dicarboxylate* **2 (i)**. Synthesized used general procedure for step (b). Yield 80%, clear colorless oil. ^1^H NMR (Chloroform-*d*, 400 MHz) δ 4.27 (1H, h, *J*=5.7 Hz), 4.11 (1H, ddd, *J*=28.2, 8.8, 5.6 Hz), 3.62 (1H, ddd, *J*=38.5, 10.9, 6.1 Hz), 3.18 (1H, ddd, *J*=19.5, 10.9, 5.0 Hz), 2.31 (1H, dddd, *J*=18.9, 12.9, 8.8, 5.9 Hz), 1.91 (1H, dt, *J*=13.0, 5.5 Hz), 1.46 – 1.35 (18H, m), 0.82 (9H, d, *J*=4.4 Hz), 0.00 (6H, s). ^13^C NMR (Chloroform-*d*, 101 MHz) δ 171.31, 153.90, 80.84, 79.77, 69.77, 58.34, 53.98, 39.45, 28.41, 28.11, 25.84, 18.17.

*(2R,4R)-di-tert-butyl 4-((tert-butyldimethylsilyl)oxy)pyrrolidine-1,2-dicarboxylate* **2 (ii)**. Compound was synthesized as 2 (ii) above following step (b) general procedure. Yield 86%, clear colorless oil. ^1^H NMR (Chloroform-*d*, 400 MHz) δ 4.29 (1H, h, *J*=5.6 Hz), 4.13 (1H, ddd, *J*=28.5, 8.8, 5.6 Hz), 3.64 (1H, ddd, *J*=39.5, 10.9, 6.1 Hz), 3.21 (1H, ddd, *J*=19.4, 10.9, 5.1 Hz), 2.44 – 2.23 (1H, m), 1.93 (1H, dt, *J*=13.1, 5.5 Hz), 1.51 – 1.37 (18H, m), 0.85 (9H, d, *J*=4.3 Hz), 0.03 (6H, s). ^13^C NMR (Chloroform-*d*, 101 MHz) δ 171.38, 153.96, 80.92, 79.85, 69.81, 58.40, 54.01, 39.50, 28.46, 28.16, 25.89, 18.23

*(2R,4S)-di-tert-butyl 4-((tert-butyldimethylsilyl)oxy)pyrrolidine-1,2-dicarboxylate* **2 (iii)**. Compound was synthesized following step (b) general procedure. Yield 90%, clear colorless oil. ^1^H NMR (Chloroform-*d*, 400 MHz) δ 4.40 (1H, p, *J*=5.0 Hz), 4.23 (1H, ddd, *J*=31.4, 8.3, 6.0 Hz), 3.57 (1H, ddd, *J*=16.5, 10.9, 5.2 Hz), 3.42 – 3.17 (1H, m), 2.24 – 2.06 (1H, m), 1.99 (1H, dt, *J*=12.8, 5.4 Hz), 1.53 – 1.38 (18H, m), 0.86 (9H, s), 0.05 (6H, d, *J*=1.7 Hz). ^13^C NMR (Chloroform-*d*, 101 MHz) δ 172.26, 154.06, 80.94, 79.83, 69.57, 58.75, 54.27, 39.79, 28.34, 28.01, 25.72, 18.02.

*(2S,4S)-di-tert-butyl 4-hydroxypyrrolidine-1,2-dicarboxylate* **3 (i)**. Synthesized according to step (c) general procedure. Yield 99%, white solid formed by trituration of colorless oil residue in DCM and hexanes. ^1^H NMR (Chloroform-*d*, 400 MHz) δ 4.27 (1H, dt, *J*=9.5, 4.5 Hz), 4.15 (1H, ddd, *J*=21.1, 9.8, 1.8 Hz), 3.78 – 3.41 (3H, m), 2.26 (1H, dddd, *J*=20.4, 14.3, 9.8, 4.8 Hz), 2.10 – 1.94 (1H, m), 1.44 (9H, d, *J*=2.5 Hz), 1.41 (9H, d, *J*=4.8 Hz). ^13^C NMR (Chloroform-*d*, 101 MHz) δ 174.21, 153.94, 82.30, 80.29, 70.30, 58.83, 55.65, 38.77, 28.41, 27.97.

*(2R,4R)-di-tert-butyl 4-hydroxypyrrolidine-1,2-dicarboxylate* **3 (ii)**. Synthesized as compound 3 (ii) above. Yield 99%, white solid. ^1^H NMR (Chloroform-*d*, 400 MHz) δ 4.29 (1H, t, *J*=4.4 Hz), 4.25 – 4.02 (1H, m), 3.78 – 3.34 (3H, m), 2.27 (1H, dddd, *J*=20.2, 14.4, 10.5, 4.7 Hz), 2.08 – 1.92 (1H, m), 1.47 (9H, d, *J*=2.2 Hz), 1.43 (9H, d, *J*=4.2 Hz). ^13^C NMR (Chloroform-*d*, 101 MHz) δ 173.26, 152.87, 81.31, 79.26, 69.33, 57.79, 54.69, 37.71, 27.35, 26.92.

*(2R,4S)-di-tert-butyl 4-hydroxypyrrolidine-1,2-dicarboxylate* **3 (iii)**. Compound synthesized according to general procedure for step (c). Yield, quantitative (100%). Colorless oil which turned solid on freezing. ^1^H NMR (Chloroform-*d*, 400 MHz) δ 4.28 (1H, p, *J*=3.8 Hz), 4.13 (1H, q, *J*=7.6, 7.1 Hz), 4.02 (1H, s), 3.49 – 3.21 (2H, m), 2.26 – 2.00 (1H, m), 1.86 (1H, ddd, *J*=12.8, 7.2, 5.0 Hz), 1.39 – 1.22 (18H, m). ^13^C NMR (Chloroform-*d*, 101 MHz) δ 172.19, 154.20, 80.96, 80.02, 68.67, 58.52, 54.38, 38.96, 28.20, 27.85.

*(2S,4S)-di-tert-butyl 4-(([1,1’-biphenyl]-4-carbonyl)oxy)pyrrolidine-1,2-dicarboxylate* **4 (i)**. Synthesized according to Steglich esterification general step (d) procedure. Yield 90%, white solid. ^1^H NMR (Chloroform-*d*, 400 MHz) δ 8.07 (2H, dd, *J*=8.3, 5.1 Hz), 7.61 (4H, td, *J*=8.3, 7.4, 2.0 Hz), 7.44 (2H, t, *J*=7.5 Hz), 7.38 (1H, t), 5.52 (1H, ddt, *J*=7.6, 5.7, 2.3 Hz), 4.38 (1H, ddd, *J*=47.5, 9.6, 2.1 Hz), 3.93 – 3.62 (2H, m), 2.56 (1H, dddd, *J*=20.4, 14.8, 9.7, 5.4 Hz), 2.46 – 2.31 (1H, m), 1.47 (9H, d, *J*=8.0 Hz), 1.41 (9H, d, *J*=3.8 Hz). ^13^C NMR (Chloroform-*d*, 101 MHz) δ 170.76, 165.92, 153.90, 145.91, 139.99, 130.44, 128.98, 128.52, 128.23, 127.32, 126.98, 81.20, 80.09, 72.48, 58.52, 52.36, 36.75, 28.40, 28.08.

*(2R,4R)-di-tert-butyl 4-(([1,1’-biphenyl]-4-carbonyl)oxy)pyrrolidine-1,2-dicarboxylate* **4 (ii)**. Compound was synthesized following step (d) general procedure. Yield 91%, white solid. ^1^H NMR (Chloroform-*d*, 400 MHz) δ 8.08 (2H, dd, *J*=8.2, 4.8 Hz), 7.62 (4H, td, *J*=7.9, 1.3 Hz), 7.46 (2H, t, *J*=7.5 Hz), 7.43 – 7.36 (1H, m), 5.53 (1H, ddt, *J*=7.6, 2.1 Hz), 4.39 (1H, ddd, *J*=47.8, 9.7, 2.1 Hz), 3.94 – 3.61 (2H, m), 2.57 (1H, dddd, *J*=21.1, 14.8, 9.7, 5.4 Hz), 2.48 – 2.25 (1H, m), 1.48 (9H, d, *J*=8.4 Hz), 1.41 (9H, d, *J*=4.1 Hz). ^13^C NMR (Chloroform-*d*, 101 MHz) δ 170.86, 166.05, 153.99, 146.02, 140.12, 130.53, 129.07, 128.59, 128.31, 127.42, 127.09, 81.32, 80.22, 72.55, 58.61, 52.42, 36.85, 28.49, 28.17.

*(2R,4S)-di-tert-butyl 4-(([1,1’-biphenyl]-4-carbonyl)oxy)pyrrolidine-1,2-dicarboxylate* **4 (iii)**. Synthesized using general procedure (step d). Yield 95%, white solid. ^1^H NMR (Chloroform-*d*, 400 MHz) δ 8.16 – 7.96 (2H, m), 7.73 – 7.57 (4H, m), 7.47 (2H, t, *J*=7.5 Hz), 7.40 (1H, d, *J*=7.3 Hz), 5.52 (1H, tt, *J*=5.0, 2.7 Hz), 4.38 (1H, dt, *J*=32.0, 7.7 Hz), 3.95 – 3.54 (2H, m), 2.68 – 2.42 (1H, m), 2.41 – 2.19 (1H, m), 1.48 (18H, d, *J*=8.1 Hz). ^13^C NMR (Chloroform-*d*, 101 MHz) δ 171.75, 165.95, 154.04, 146.13, 140.00, 130.32, 129.07, 128.58, 128.36, 127.40, 127.22 (d, *J*=2.4 Hz), 81.59 (d, *J*=3.6 Hz), 80.49, 72.68, 58.73 (d, *J*=3.4 Hz), 52.18, 36.97, 28.46, 28.15.

*(2S,4R)-di-tert-butyl 4-(([1,1’-biphenyl]-4-carbonyl)oxy)pyrrolidine-1,2-dicarboxylate* **4 (iv)**. Synthesized according to scheme 2 step (d) general procedure. Yield 90%. Yellow solid. ^1^H NMR (Chloroform-*d*, 600 MHz) δ 8.10 – 8.04 (2H, m), 7.65 (2H, dd, *J*=8.2, 3.4 Hz), 7.63 – 7.58 (2H, m), 7.48 – 7.43 (2H, m), 7.42 – 7.36 (1H, m), 5.52 (1H, ddt, *J*=7.7, 2.4 Hz), 4.38 (1H, dt, *J*=49.0, 7.7 Hz), 3.91 – 3.64 (2H, m), 2.62 – 2.45 (1H, m), 2.31 (1H, dddd, *J*=15.3, 10.2, 7.4, 5.5 Hz), 1.48 (18H, d, *J*=12.3 Hz). 13C NMR (Chloroform-d, 151 MHz) δ 171.72, 165.90, 154.02, 146.09, 139.94 (d, J=5.5 Hz), 130.28, 129.03, 128.55, 128.32, 127.36, 127.16, 81.54, 80.47, 72.65, 58.68, 52.15, 36.93, 28.43, 28.12.

*(2S,4S)-4-(([1,1’-biphenyl]-4-carbonyl)oxy)pyrrolidine-2-carboxylic acid* (*Lc*-BPE) **5 (i)**, *(2R,4R)-4-(([1,1’-biphenyl]-4-carbonyl)oxy)pyrrolidine-2-carboxylic acid* (*Dc*-BPE) **5 (ii)**, (*2R,4S)-4-(([1,1’-biphenyl]-4-carbonyl)oxy)pyrrolidine-2-carboxylic acid* (*Dt*-BPE) **5 (iii)** and *(2S,4R)-4-(([1,1’-biphenyl]-4-carbonyl)oxy)pyrrolidine-2-carboxylic acid* (*Lt*-BPE) **5 (iv)** were prepared according to general procedure for step (e) and precipitated from methanol with diethyl ether as white powder (TFA salts). Yields, 60%. 71%, 82% and 90% respectively. The final product was confirmed by TLC (20 - 40% methanol in DCM), rf ∼ 0.2) and used as it is in all experiments without further purification.

## Acknowledgements

This study was supported by a grant from the National Institutes of Health (http://www.nih.gov.eresources.mssm.edu) (R01 GM108911) to A.S. and C.G. and T32 CA078207 to R.A.G., the National Science Foundation Grant 1515028 awarded to C.G, the NWO TOP grant 714.018.003 to D.J.S; and the NWO Veni grant 722.017.001 and NWO Start-Up grant 740.018.016 to C.P. M.B. would like to acknowledge the INCEPTION project ANR-16-CONV-0005 and PRACE for awarding access to Piz Daint at CSCS, Switzerland. We also thank J. Rheinberger for help with cryo-EM data acquisition; M. Punterfor maintenance of the image-processing cluster; V. Arkhipova for help in MSP2N2 purification; T. Postmus for initial tests in proteoliposomes.

## Contributions

R.A.G., E.N., A.G., M.B., D.J.S., C.P., C.G., A.S. conceptualized and designed the study. Homology modeling and molecular docking was done by R.A.G., supervised by A.S. Compound synthesis and biochemical experiments were performed by E.N. and C.G. Cryo-EM studies, model building and proteoliposome assays were done by A.G., supervised by D.J.S. and C.P. Molecular dynamics simulations were performed by M.B. The manuscript was drafted by R.A.G. and A.S. and R.A.G., E.N., A.G., M.B., D.J.S., C.P., C.G., A.S. contributed to the writing of the final manuscript.

