## Extended_Data_Figures for "Structural basis for stereospecific inhibition of ASCT2 from rational design"

### Extended Data Fig. 1

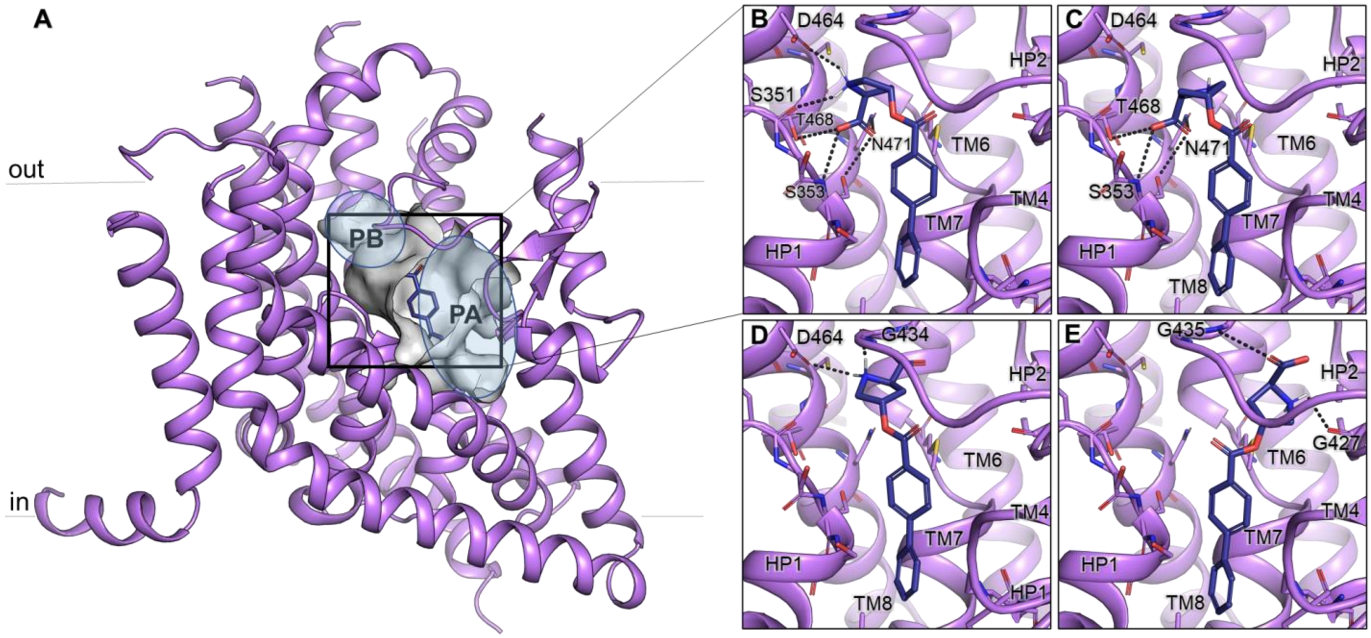


**Extended Data Fig. 1. Predicted binding mode of ASCT2 inhibitors. (A)** The outward-open homology model of ASCT2 based on the EAAT1 structure (PDB ID: 5MJU) is shown as a purple cartoon. Subpocket A and B (PA and PB, respectively) of the binding site are highlighted with ellipsoids. Key amino acids in the binding site are shown as sticks, where oxygen, nitrogen, and sulfur atoms are shown in red, blue, and yellow respectively; hydrogen bonds are represented with grey dashes. **(B-E)** The four ASCT2 inhibitors are *cis* and *trans* isomers of L and D - proline derivatives (*Lc*-BPE, *Dc*-BPE, *Dt*-BPE, *Lt*-BPE respectively) are predicted to bind a conformation-specific pocket accessible in the outward-open conformation, where HP2 is in an open position.

**Extended Data Fig. 2**

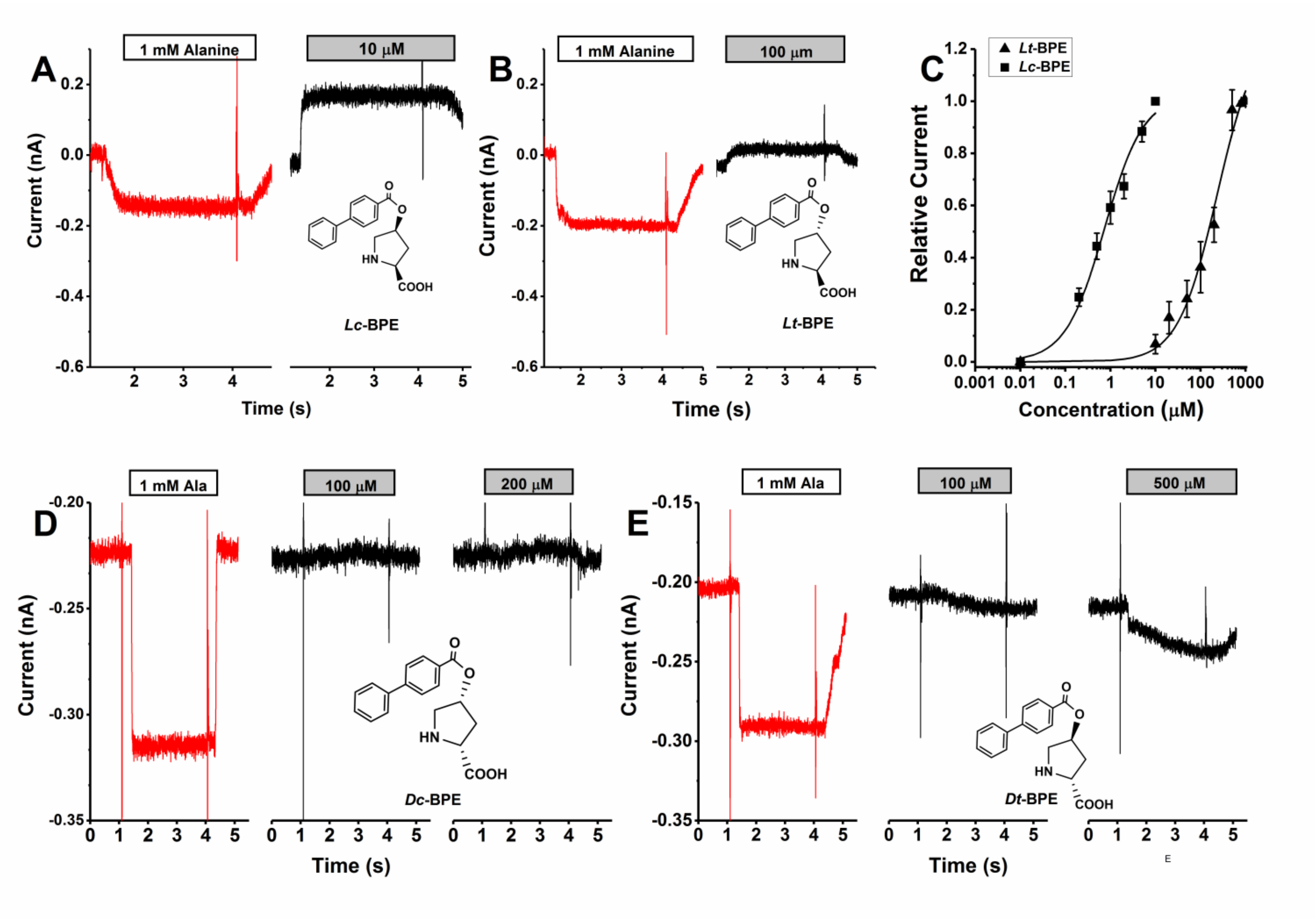


**Extended Data Fig. 2: Electrophysiological characterization of 4-hydroxyproline biphenyl ester diastereomers. (A & B)** Original current traces of inward anion currents produced by the application of 1 mM alanine (red trace) and inhibition of leak anion current by the application of *Lc*-BPE (A) and *Lt*-BPE (B) to rASCT2 expressing cells. The white and grey bars illustrate the time of compound application to rASCT2. **(C)** Dose response curves for *Lc*-BPE (squares) and *Lt*-BPE (triangles). The solid lines represent fits according to Michaelis-Menten-like equation with $K_i$ of 0.74 ± 0.11 μM and 232 ± 44 μM. **(D)** and **(E)** Analogous experiment as in **(A)** and **(B)** but D-isomers of 4-hydroxyproline, *Dc*-BPE **(D)** and *Dt*-BPE **(E)** were used. Currents from these two isomers were either too small at high compound concentrations (*Dc*-BPE) or unspecific (*Dt*-BPE) and therefore dose response curves and estimated $K_i$ could not be obtained. Transmembrane potential was 0 mV in the presence of 130 mM NaSCN/10 mM Alanine internal, and 140 mM NaCl external.

**Extended Data Fig. 3**

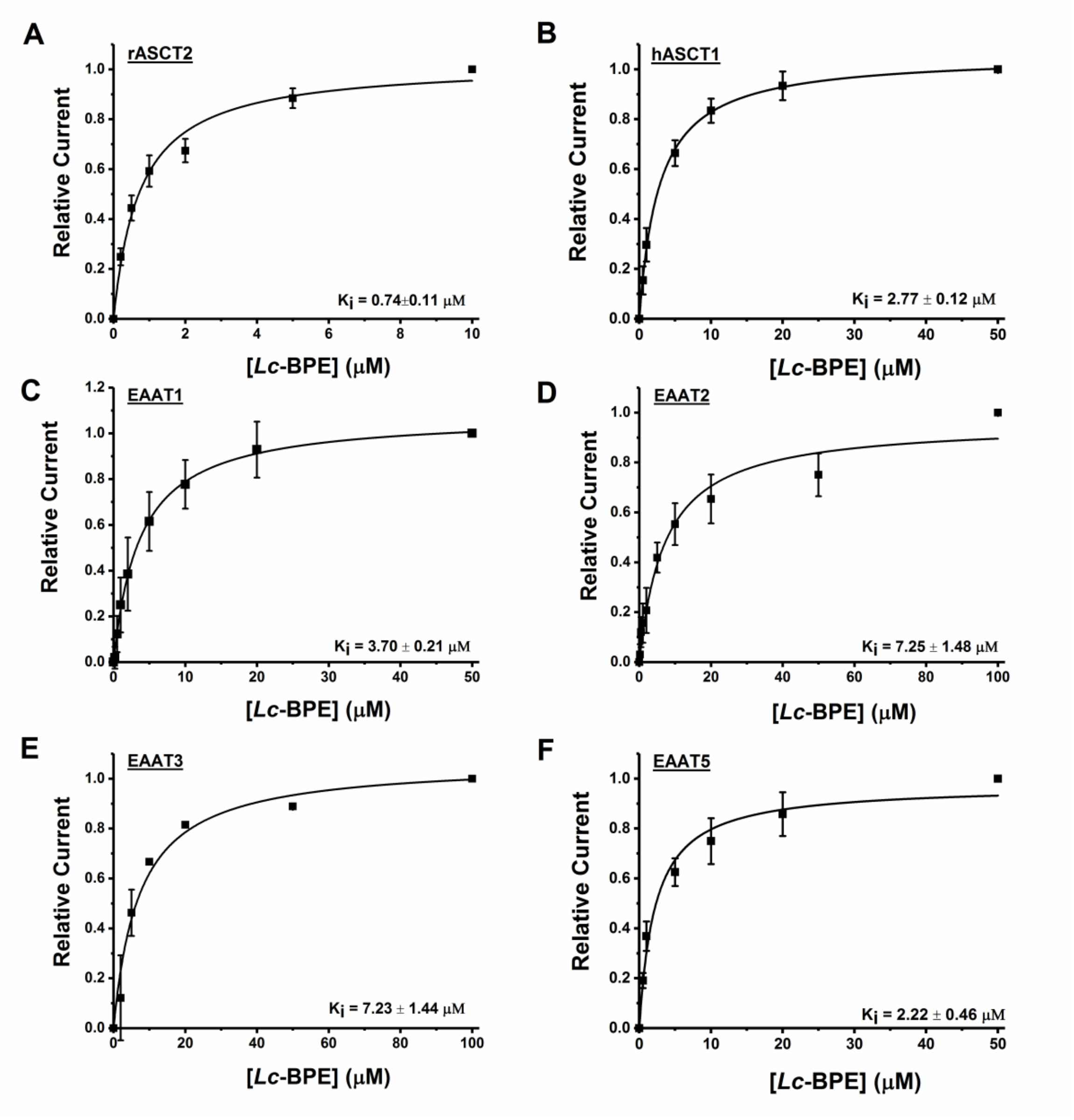


**Extended Data Fig. 3. SLC-1 specificity of *Lc*-BPE. (A-F)** Dose response curves obtained from currents recorded after application of *Lc*-BPE extracellularly to cells transiently expressing rASCT2 (as shown in Fig. 2B), hASCT1, rEAAT1, rEAAT2, rEAAT3 and hEAAT5 respectively. For rASCT2 and hASCT1, intracellular pipette solution contained 10 mM alanine and 130 mM NaSCN. For rEAAT1, rEAAT2, rEAAT3 and hEAAT5, intracellular solution contained 10 mM glutamate and 130 mM NaSCN. Extracellular solution was the same for all the experiments (140 mM NaCl). The voltage was 0 mV.

### Extended Data Fig. 4

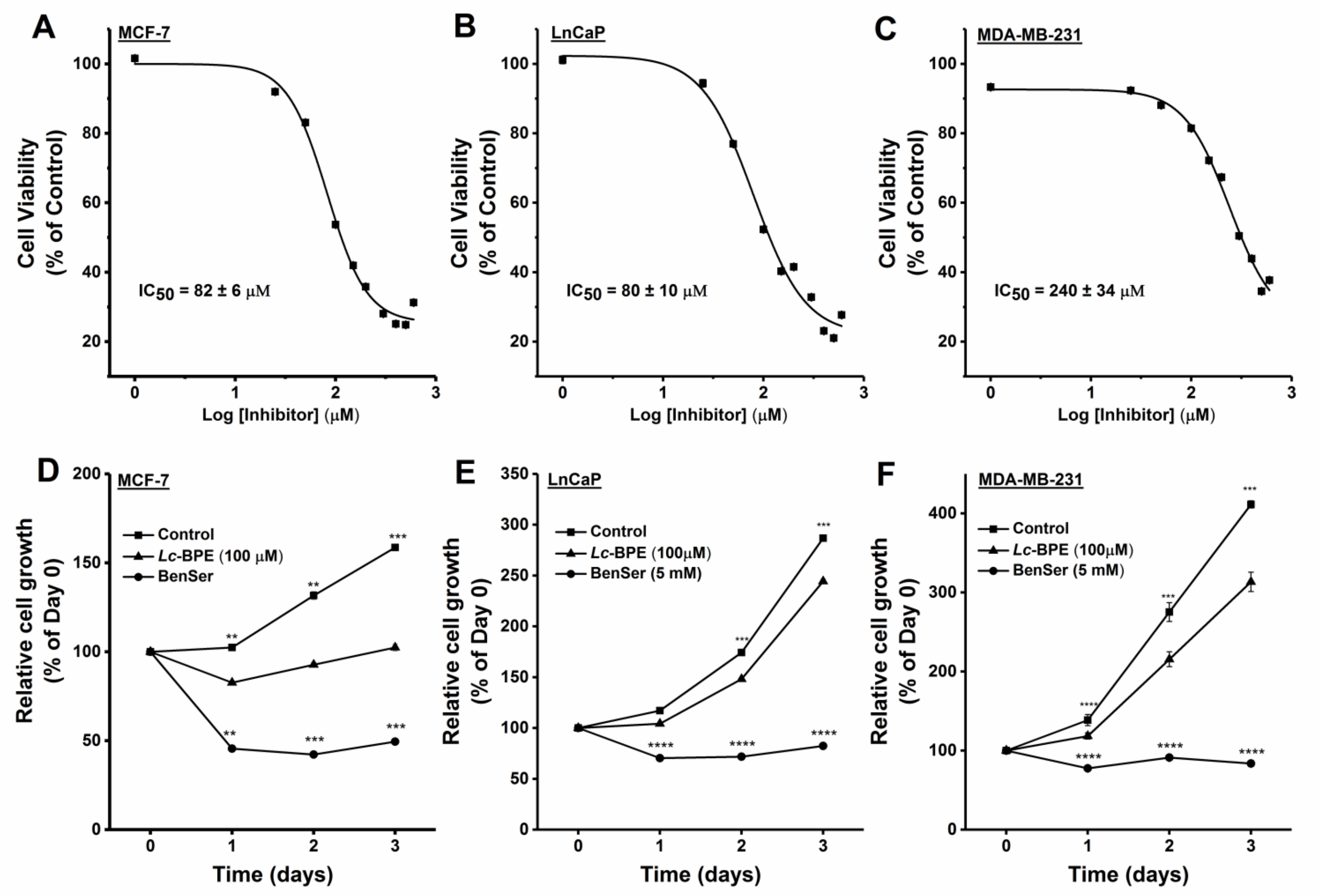


**Extended Data Fig. 4. *Lc*-BPE inhibits proliferation in LnCaP, MCF-7 and MDA-MB-231 cancer cell lines. in a dose dependent manner**. **(A-C)** Dose response graphs of human cancer cell lines **(A)** MCF-7, **(B)** LnCap and **(C)** MBA-MB-231. Absorbance from different *Lc*-BPE inhibitor concentrations was normalized to the absorbance at 1 µM *Lc*-BPE inhibitor. **(D-E)** Relative cell viability in absence (squares) or presence of *Lc*-BPE inhibitor (triangles) and benzyl serine (BenSer, circles) measured by MTT assay in **(D)** MCF-7, **(E)** LnCaP , and **(F)** MDA-MB-231 cancer cells. Absorbance for each day was normalized to absorbance for Day 0 (before treatment with *Lc*-BPE or BenSer).

**Extended Data Fig. 5**

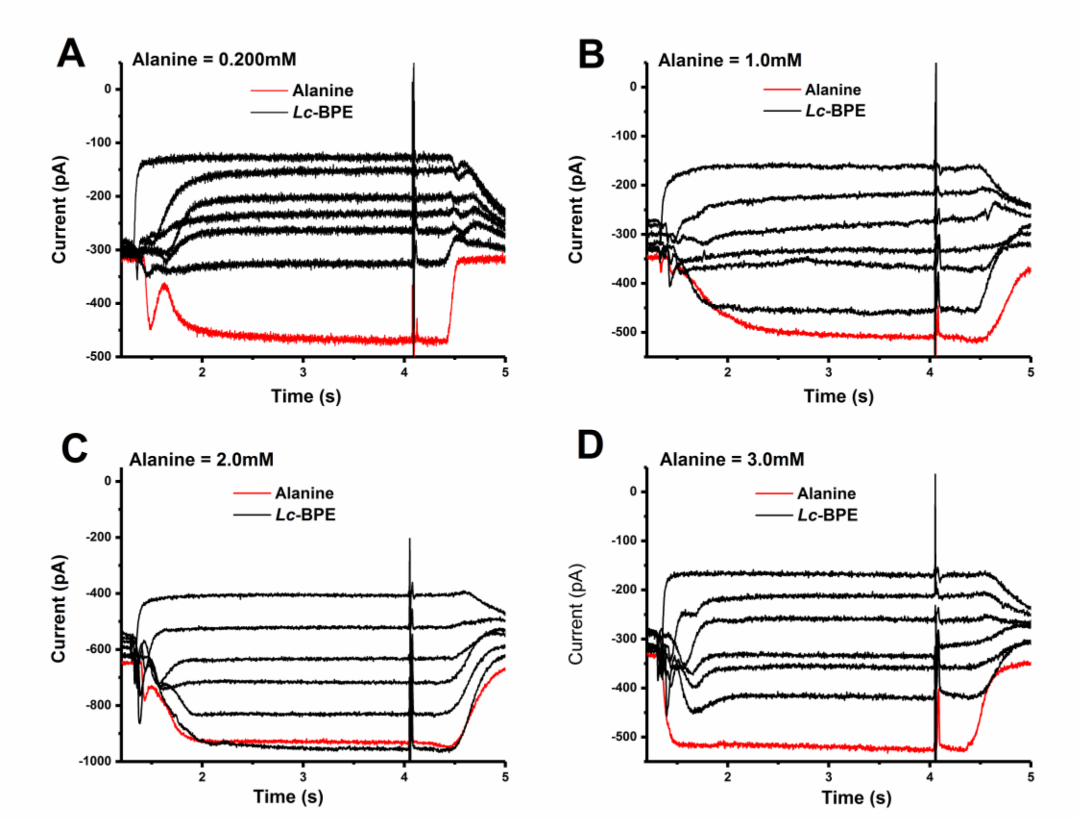


**Extended Data Fig. 5. *Lc*-BPE-induced currents in the presence of varying concentrations of alanine.** Currents were produced by the application of different concentrations of alanine in the absence (red trace) and presence (black traces) of varying *Lc*-BPE concentrations to HEK293T cells overexpressing rASCT2. (**A)** 0.2 mM alanine, **(B)** 1.0 mM Alanine, (**C)** 2.0 mM alanine and (**D)** 3.0 mM alanine. All experiments were done at 0 mV, intracellular and extracellular buffers consisted of 130 mM NaSCN and 140 mM NaCl respectively at pH 7.40. Both alanine and *Lc*-BPE were dissolved in extracellular buffer and applied externally.

**Extended Data Fig. 6**

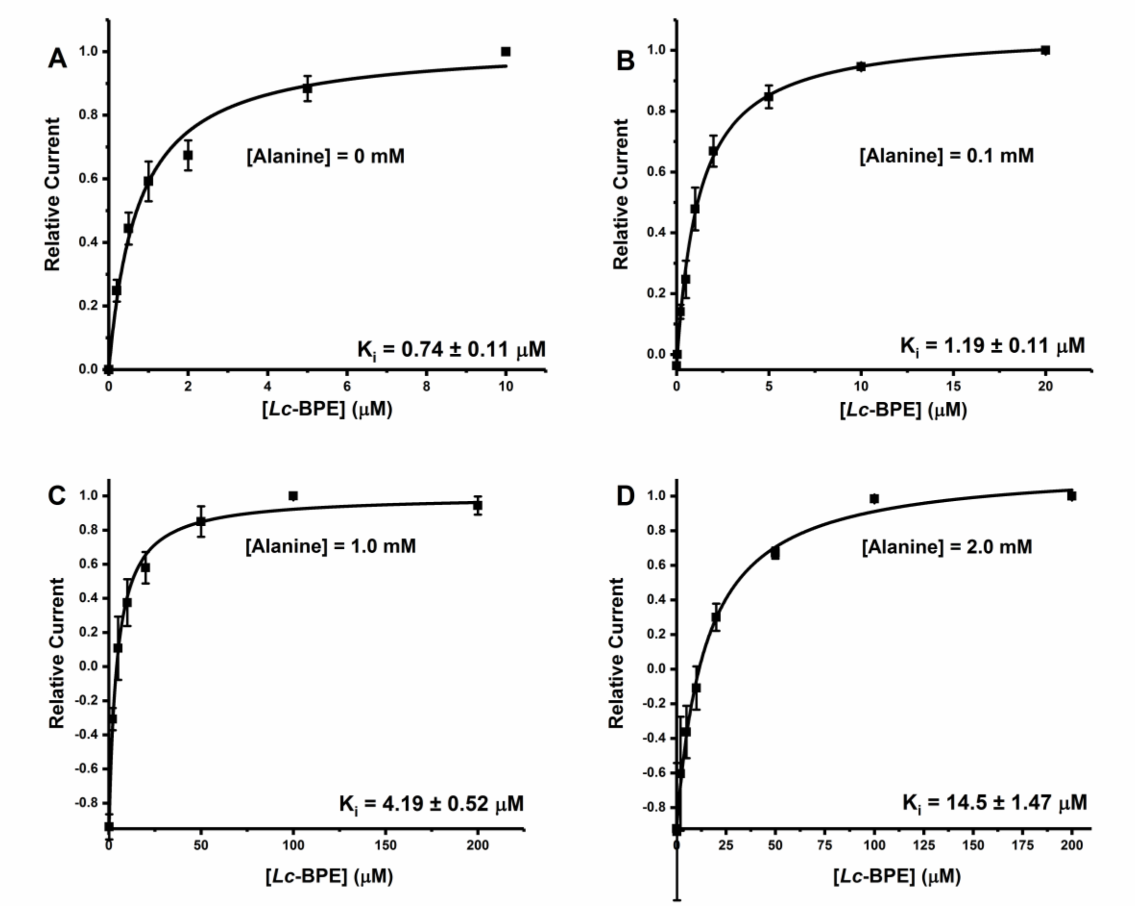


**Extended Data Figure 6: *Lc*-BPE dose response relationships in the presence of varying concentrations of alanine. (A)**, no alanine, **(B)**, 0.1 mM alanine, **(C)**, 1.0 mM Alanine and **(D)**, 2.0 mM alanine. The lines represent fits of the currents according to the equation: $I = I_1 + I_2$ [Inh]/($K_i$ + [Inh]), where $I_1$ is the alanine induced current without the inhibitor, and $I_2$ is the maximum current in the presence of saturating inhibitor concentration, $[Inh]_{max}$[1]. For **(A)** and **(B)**, currents are normalized to the current recorded after application of highest inhibitor concentration to rASCT2 expressing cells while **C** and **D** currents are normalized to the current recorded after application of alanine in the absence of inhibitor (those two approached results to the same $K_i$).

**Extended Data Fig. 7**

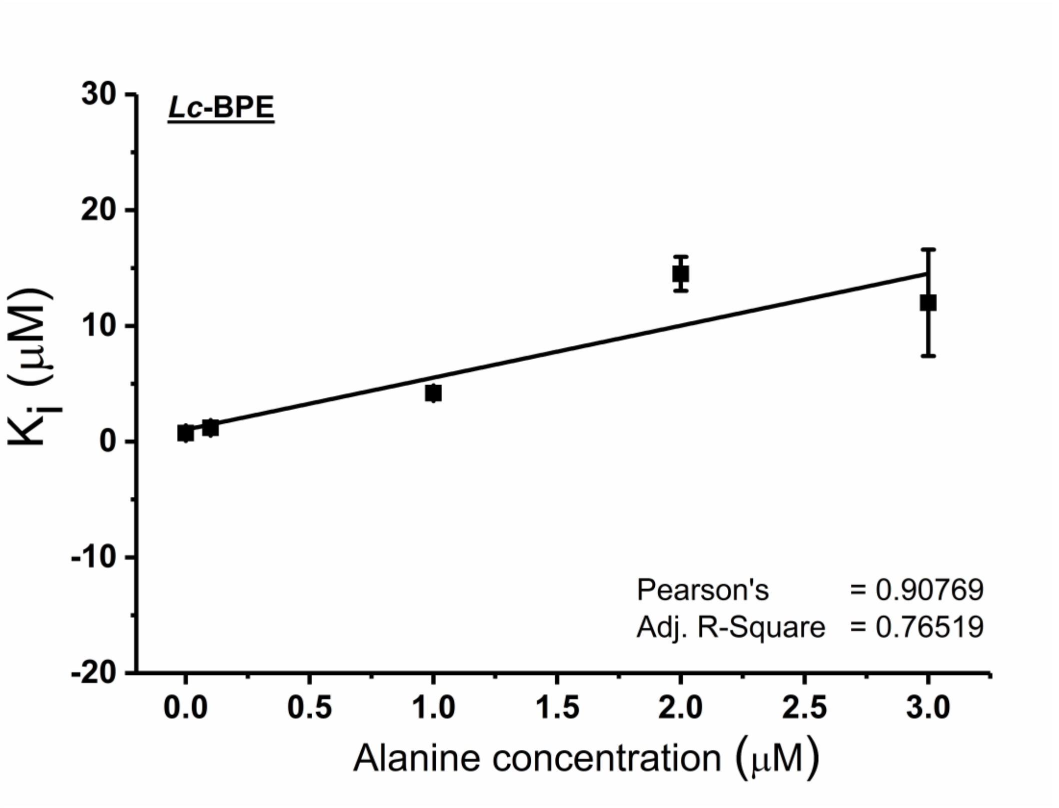


**Extended Data Figure 7: *Lc*-BPE competes with substrate for the substrate binding site.** *Lc*-BPE apparent affinity ($K_i$) (obtained from Extended Data Fig. 6) plotted as a function of alanine concentration, exhibiting a linear relationship according to the equation $K_i(Ala) = K_i(0) + [Ala]K_i(0)/K_m(Alanine)$, $K_i(Ala)$ and $K_i(0)$ are $K_i$ values in the presence and absence of alanine while $K_m(Alanine)$ is the apparent Michaelis-Menten constant for alanine activation of inwardly directed anion current[2] . Pearson's r value is 0.91 and adjusted $R^2$ is 0.77 which demonstrates good correlation.

**Extended Data Fig. 8**

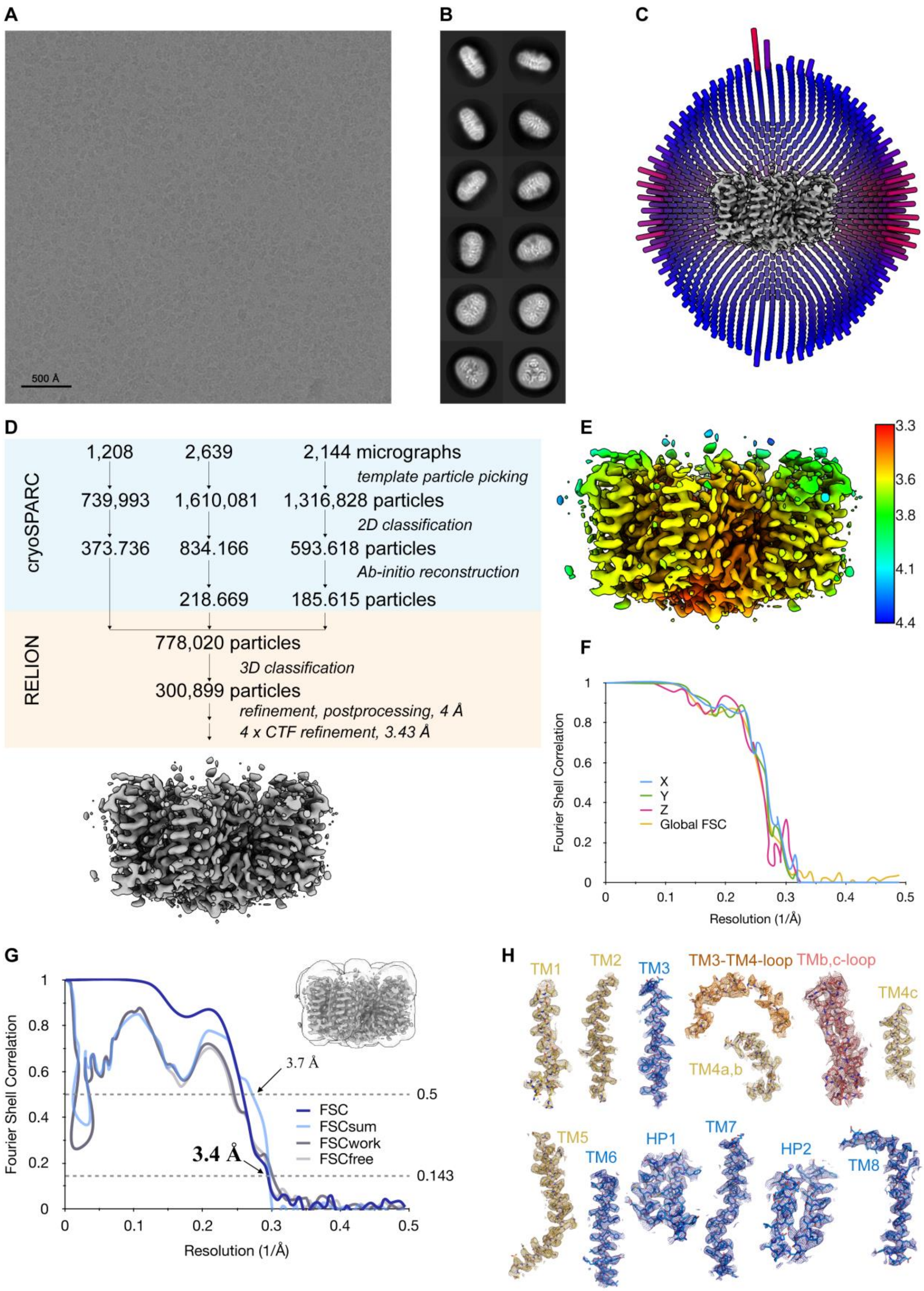

**Extended Data Fig. 8. Cryo-EM reconstruction of ASCT2 nanodiscs in the presence of inhibitor.** Representative cryo-EM image **(A)** and 2D-class averages **(B)** of vitrified ASCT2 nanodiscs in the presence of inhibitor. **(C)** Angular distribution plot of particles included in the final C3-symmetrized 3D reconstruction. **(D)** Image processing workflow. **(E)** Final reconstructed map coloured by local resolution, as estimated in Relion. **(F)** Anisotropy estimation plot of the final map. The global FSC curve is represented in yellow. The directional FSCs along the x, y and z axis are displayed in blue, green and red, respectively. **(G)** FSC plot used for resolution estimation and model validation. The gold-standard FSC plot between two separately refined half-maps is shown in blue and indicates a final resolution of 3.4 Å. The FSC model validation curves for FSCsum, FSCwork and FSCfree, as described in material and methods, are shown in light blue, dark grey and light grey respectively. A thumbnail of the mask used for FSC calculation overlaid on the map is shown in the upper right corner. Dashed lines indicate the FSC thresholds used for FSC of 0.143 and for FSCsum of 0.5. **(H)** Cryo-EM densities. Shown are selections of cryo-EM densities of ASCT2 nanodisc in presence of inhibitor, with the respective refined models superimposed. Models are shown as sticks and structural elements are labelled. Transmembrane helices (TM) of the transport domain are coloured in blue, of the scaffold domain in yellow, the loop between $TM4_b$ and $TM4_c$ in red, the loop between TM3 and TM4 in orange. Densities were sharpened with a b-factor of -189 Å2. Densities of TM1, TM6 and TM7 were contoured at 3 σ; TM2, TM3, TM4a,b, TM4c, TM5, TM8, HP1, HP2 were contoured at 4 σ; $TM4_{b,c}$ loop, TM3-TM4 loop was contoured at 2 σ.

**Extended Data Figure 9**

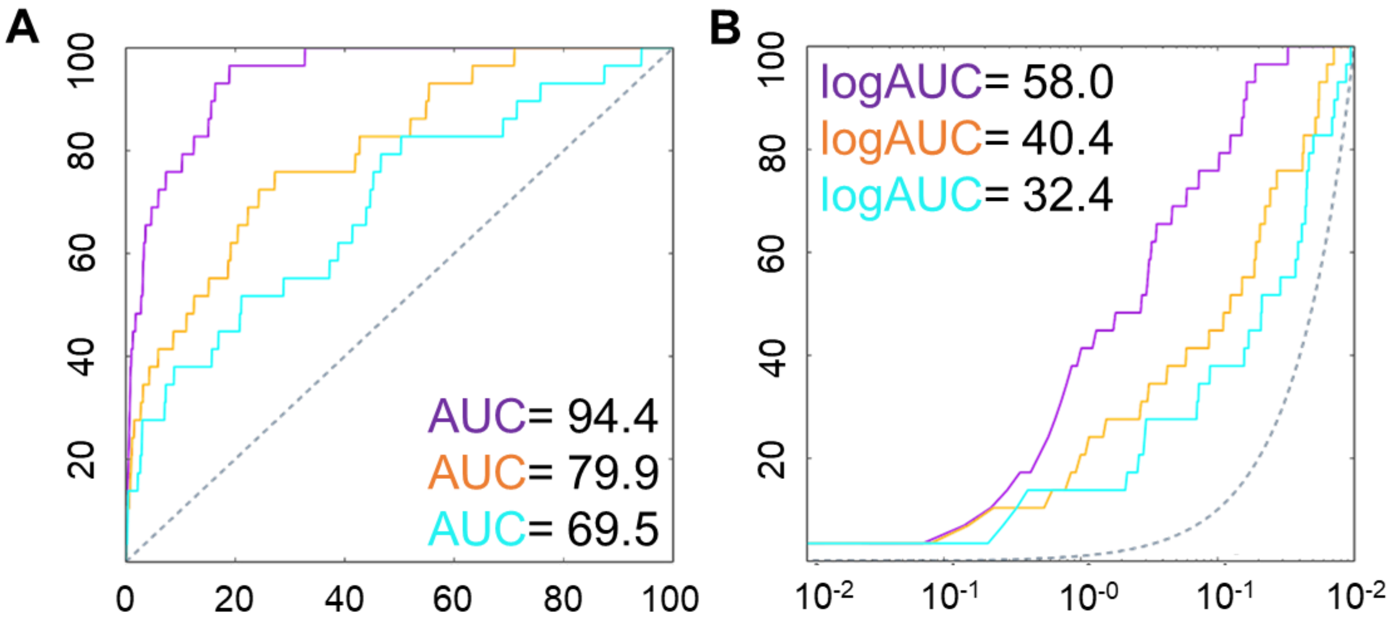


**Extended Data Figure 9. Model and structure evaluation with enrichment plots. (A) and (B)** Enrichment plots of ASCT2 homology model (purple) and structures in “ligand down” conformation before (cyan) and after sidechain refinement (orange). **(A)** Area under the curve. **(B)** log of the area under the curve. The plot for a random selection of ligands is represented by the gray dashed line.

**Extended Data Figure 10**

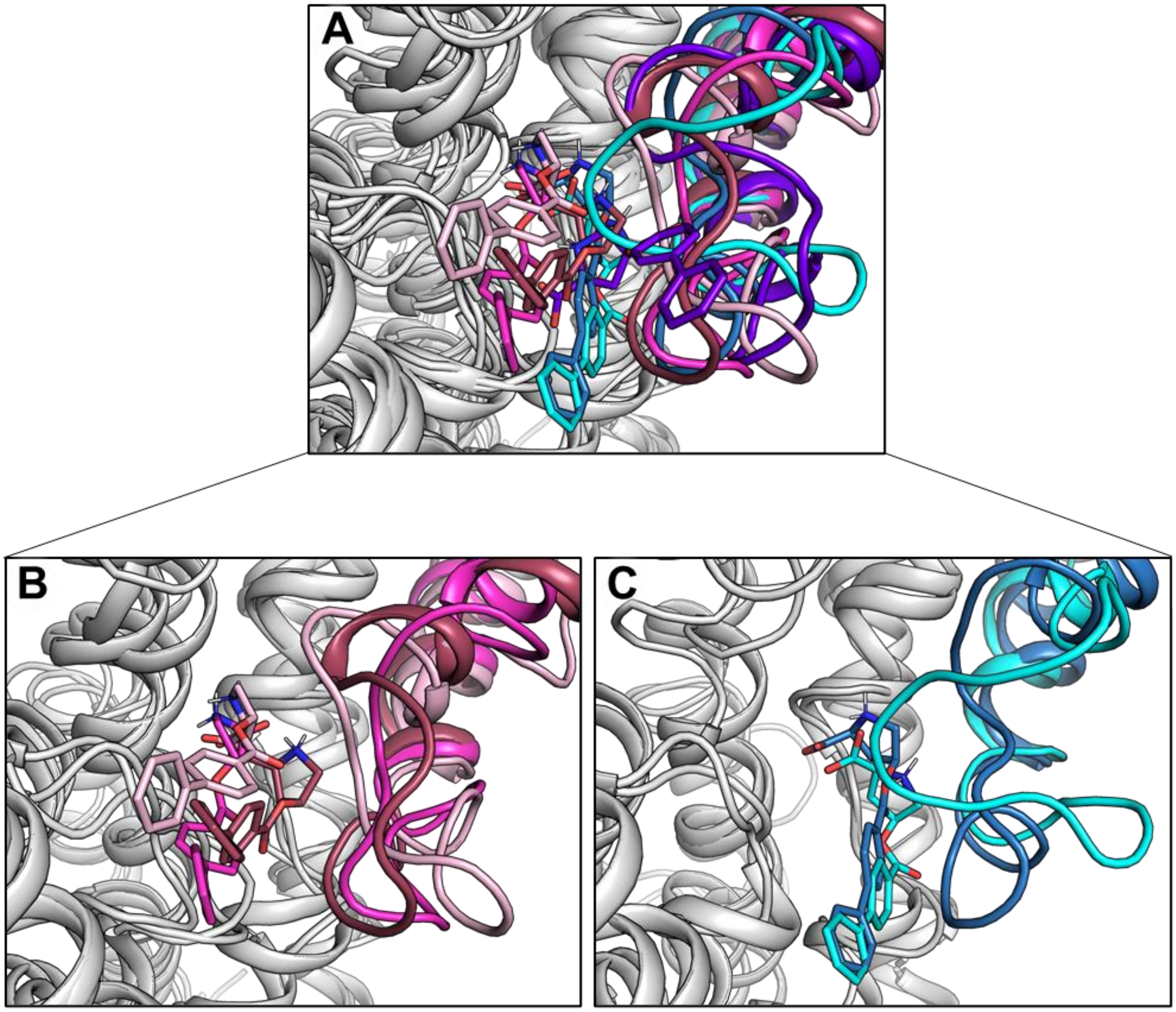


**Extended Data Figure 10. Clusters of ligand conformations obtained with metainference molecular dynamics (MD) simulations. (A)** Superposition of 6 clusters of ligand conformations in the substrate binding site. **(B)** “Ligand up” clusters in red (69%), magenta (7%) and pink (6%). **(C)** “Ligand down” clusters in dark blue and cyan represent 13% of all clusters. The purple cluster represents an outlier cluster that is outside of the binding site and is unlikely to be physiological.

**Extended Data Figure 11**

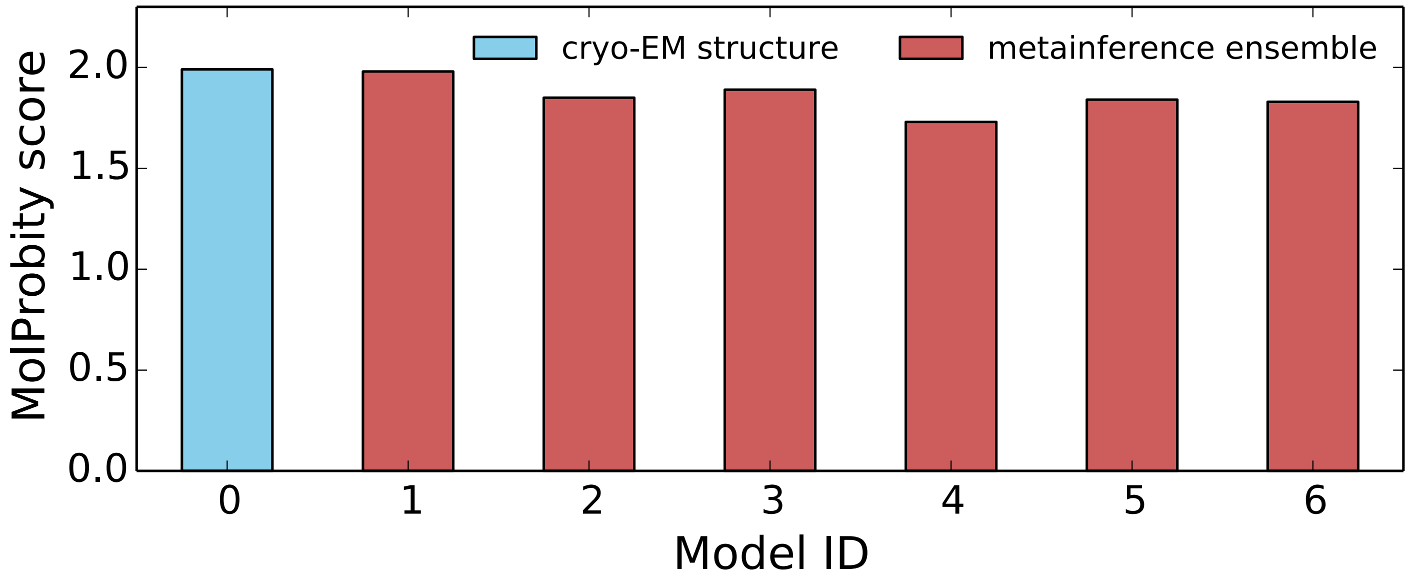


**Extended Data Figure 11. Stereochemical quality of the cryo-EM structure and metainference ensemble.** We assessed the stereochemical quality of the cryo-EM structure (cyan, model 0) as well as of the six representative clusters from the metainference ensemble (red, models 1-6) using the MolProbity score[3]. This score provides a global measure of the quality of the models in terms of number of steric clashes, rotamer outliers, and percentage of backbone Ramachandran conformations outside favored regions. The lower the score, the better the quality of the models.
